# Shape Factor Analysis as a Quantitative Framework for Assessing Spheroid and Organoid Morphology and Invasiveness

**DOI:** 10.64898/2026.03.10.710632

**Authors:** Brittany E. Schutrum, Jenny Deng, Ju Hee Kim, Amalie Gao, Emily Hur, Jack C. Crowley, Lu Ling, Matalin G. Pirtz, Coulter Q. Ralston, Alexander Yu. Nikitin, Claudia Fischbach

## Abstract

Morphological changes of spheroids and organoids are widely used as *in vitro* indicators of healthy and diseased tissue functions; however, quantitative methods to classify spheroid and organoid morphology are limited. In clinical breast imaging, radiologists use tumor shape as a prognostic marker, with irregular margins associated with invasive disease and increased malignancy. Here, we adapted this approach for translational research and developed a custom MATLAB algorithm to quantify the variance in radial lengths of invasive protrusions in spheroids and organoids. First, we analyzed digital phantoms by both ImageJ/FIJI shape descriptors and our radial length analysis to evaluate the capabilities of each measurement method. Subsequently, we performed the same comparisons with images from experimental spheroid and organoid datasets. We demonstrate that multivariate shape factor analysis, including radial length analysis, enables more reliable and comprehensive quantification of spheroid and organoid morphologies than standard shape descriptors alone. By enabling numerical morphological readouts, shape factor analysis can enhance phenotypic profiling of spheroids and organoids and provide valuable metrics for *in vitro* studies including high-throughput and drug screening workflows.

## Introduction

Advanced 3D model systems have emerged as critical tools to recapitulate the complex microenvironment of normal and malignant tissues in a controlled experimental environment^1–3^. Breast cancer progression is routinely studied with spheroids or organoids embedded into extracellular matrix mimetics^4–7^. Using these models, malignant progression can be estimated by analyzing time-dependent changes in spheroid/organoid shape, size, and margins using microscopy images. However, tools to objectively quantify spheroid and organoid morphology are lacking, making interpretation across experiments challenging. Given the emerging interest in spheroids^8^ and organoids^9^ for precision medicine and drug discovery, there is a widespread need for accurate and automated image analysis of their morphologies that is adaptable for data collection in high-throughput settings. The development of such techniques may benefit from approaches currently used in the clinic.

Routine imaging-based screening has contributed to making breast cancer the most diagnosed cancer in women worldwide^10^. Although primary tumors can be treated relatively effectively, invasive cancer is associated with worse clinical prognosis due to its increased likeliness to metastasize to distant organs^11^. Therefore, much emphasis has been placed on detecting cancer cell invasion early and developing therapies to interfere with this process^11^. During screening, irregular tumor margins are a primary focus for radiologists’ qualitative evaluation, as they are indicative of local invasion and thus, represent a key prognostic indicator of malignancy^12–14^. To improve diagnostic accuracy, radiologist readings are complemented by computer aided detection software^15^ that uses machine learning-based technology to highlight suspicious regions for radiologist review. Inspired by the value of computationally assistive tools in the clinic, we sought to develop a similar approach to quantify the morphology and invasion of 3D-cultured breast cancer spheroids in a research setting.

Previous approaches to classify *in vitro* spheroid invasion range from simple analysis of spheroid size^4^ to advanced machine learning algorithms^16,17^. The most straightforward methods measure the largest invasion distance from the core of a spheroid or quantify the change in area over time^4,18^. While these metrics can classify the presence of invasion, they cannot differentiate a growing, non-invasive spheroid from an invasive spheroid. With high resolution imaging single cell invasion can be quantified by segmenting the core of the spheroid and measuring the invasion distance of individual cells ^19^. Similarly, collective invasion analyses have focused on measuring the number and dimensions of collective invasion strands^20–22^. However, these collective and single cell analysis methods rely on acquiring time consuming high-resolution images and often additional staining, processes which are impractical in high-throughput settings. Characterizing a basic shape factor such as circularity from lower resolution images of spheroids/organoids has been implemented to circumvent these challenges^23^. Yet this approach also comes with limitations as the frequently used shape descriptors in ImageJ/FIJI^24^ cannot reliably differentiate invasions from other morphological features, including elongation, asymmetric invasion, or regions of concavity. Advances in machine learning can address these limitations^17^, but widespread adoption is still limited due to the need for specialized software and knowledge. Hence, there exists an unmet need for simple, user-friendly approaches to quantify the invasion or morphology of spheroids and organoids in both research and high-throughput settings.

Here, we developed a custom MATLAB code for radial length analysis of spheroid and organoid morphology using low resolution images. Our analysis pipeline is inspired by a computational approach that was initially developed to identify perimeter undulations or surface irregularities of tumors in clinical images^25^. It is designed to quantify variations in perimeter contours, represented as radial lengths, in a size agnostic manner, thus allowing for comparisons across different length scales. For shape factor quantification and comparisons, we use custom designed digital phantoms and experimental data sets. While radial length analysis is superior for specifically detecting collective radial invasion in spheroids as compared to the FIJI shape descriptors^24^, circularity or convexity is better suited to characterize organoid malignancies that manifest as folds or protrusions. These results suggest that easy-to-implement multivariate shape factor analysis is required for reliable and comprehensive quantification of spheroid and organoid morphologies using low resolution bright-field images.

## Results

### Advantages and limitations of standard shape factor analysis to quantify invasion

Shape factors are dimensionless numbers that quantify object shape using parameters such as perimeter, area, convex area, convex perimeter, major axis, and minor axis^26^ (**fig 1A**). Because they are size-independent, shape factors can compare structures across different length scales. For example, the widely used image-analysis software FIJI^24^ computes the following shape factors (termed shape descriptors in FIJI): aspect ratio, roundness (inverse of aspect ratio), circularity, and solidity^27^ whose respective formulas are displayed in **fig 1B** although the nomenclature and formulas can differ for other software programs. While not included in the FIJI shape descriptors, convexity is another metric that can be obtained by dividing the convex perimeter by the perimeter^28^ using a FIJI macro (https://github.com/BrittanySchutrum/FIJI-Spheroid-Morphological-Signatures) (**fig. 1A, B**). These different metrics vary in scale: roundness, circularity, solidity, and convexity range from 0 to 1, whereas aspect ratio ranges from 1 to infinity.

**Figure 1:**
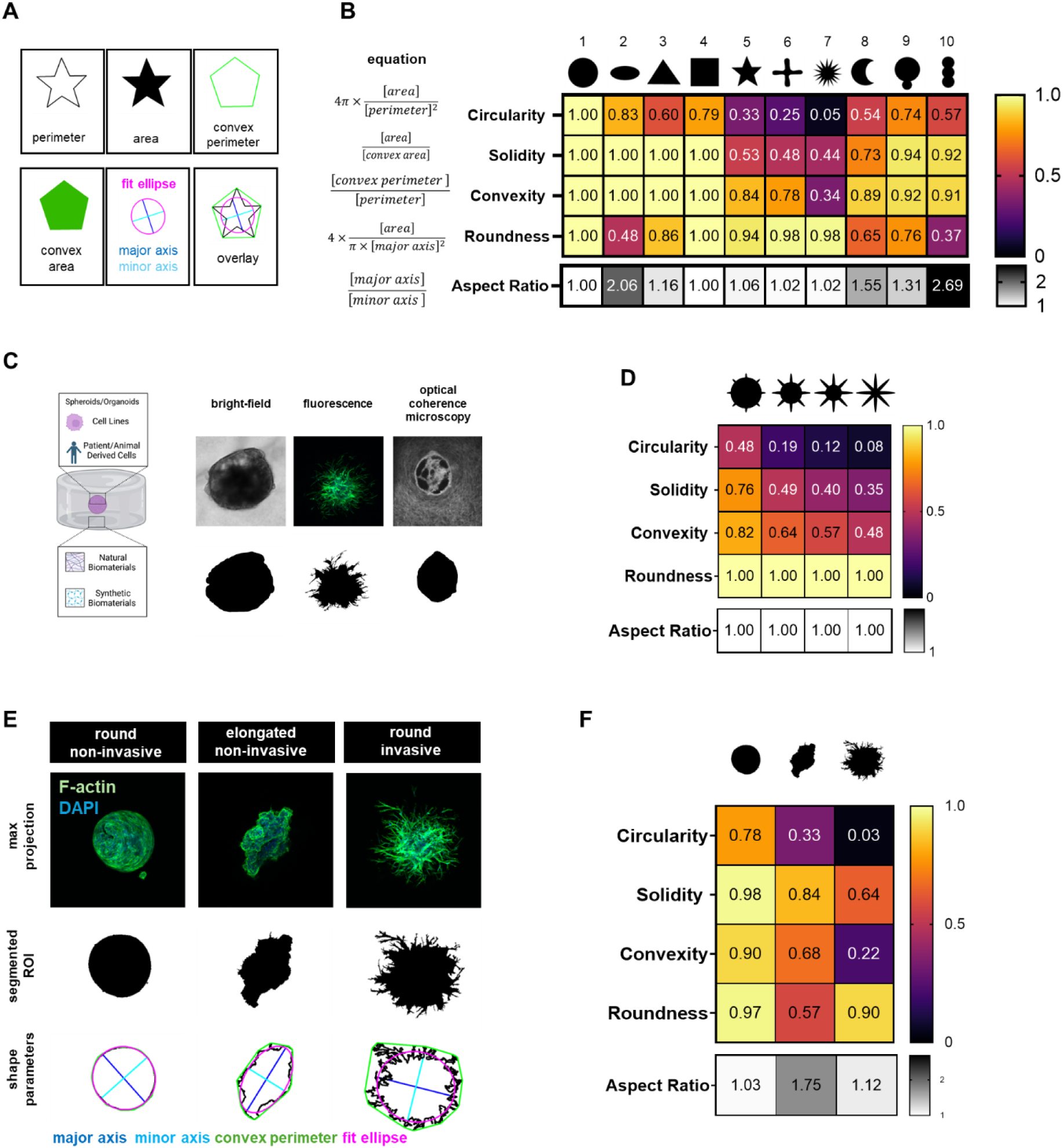
Analysis of FIJI/ImageJ shape descriptors to quantify morphological changes of digital phantoms and spheroids in experimental settings. **A)** Schematic of a star shaped digital phantom to highlight component metrics used to calculate FIJI shape descriptors. B) Heat maps visualize the capabilities of each metric to differentiate between the shapes of digital phantoms. The formula for each shape descriptor is listed to the left and each evaluated shape appears at the top of the heat map. Values were computationally determined in FIJI from regions of interest of each shape with perimeter points interpolated to a 1-pixel interval on shapes in a 512×512 resolution image. Resolution sensitivity analysis for each shape is included in supplemental information. C) Depiction of 3D invasion assays using matrix-embedded breast cancer spheroids. Microscopic images of matrix-embedded breast cancer spheroids collected with bright-field, confocal, or optical coherence microscopy and their corresponding segmented ROIs. D) Digital phantoms designed to exhibit different protrusion lengths and their characterization by different shape factors. E) Confocal maximum projections of spheroids (F-actin -green, DAPI -blue) with the thresholded and segmented ROIs displayed in black and the annotated ROIs below (major axis-blue, minor axis-cyan, convex perimeter-green, fit ellipse-magenta). F) Heat maps present the measured values of the shape descriptors for each spheroid sample in E. Schematic in C created in Biorender.com

We first designed ten digital phantoms to evaluate how each FIJI shape descriptor classifies varied morphological features. These digital phantoms were divided into three groups (**fig. 1B**): the first four (circle, oval, triangle, square) have smooth contours but vary in bulk geometry; the next three (star, X, burst) feature surface irregularities with differing numbers and sizes of protrusions; and the last three (half-moon, single bud, double bud) mimic varying convexity and budding or lobular structures. As a reference shape, the circle scores 1 for all metrics. High convexity shapes with smooth contours (circle, oval, triangle, square) all have perfect values of 1 for convexity and solidity because convex area equals area and the convex perimeter equals perimeter (**fig 1A, B**). These smooth contour shapes highlight that circularity and roundness are not interchangeable metrics: circularity depends on the ratio of area to perimeter, while roundness depends on the ratio of area to major axis (i.e. elongation). Given that a circle is the smallest perimeter for a given area, the deviation from a circle either through elongation or additional polygonal sides causes a decrease in circularity. Mathematically, the roundness of a square is 1, identical to a circle because in both cases the major and minor axes are equal. This counterintuitive result contrasts with the general use of round as a term to describe a shape with no sharp edges.

Next, we compared shapes that differ in the number and sizes of protrusions (shapes 5-7, **fig 1B**) but exhibit a roundness and aspect ratio close to 1. The convex areas of these shapes were roughly twice as large as their respective areas resulting in similar solidity values of approximately 0.5. Hence, solidity is not a robust measure to quantify shapes with varying numbers and sizes of protrusions. Instead, circularity and convexity are more appropriate metrics to describe these differences, as both are strongly influenced by the shape’s perimeter, which increases with the number of protrusions. This increase in perimeter relative to the area and convex perimeter leads to corresponding changes in circularity and convexity, respectively. Finally, we investigated alterations to a circle with a concave region, a single bud, and a double bud. As expected, roundness and aspect ratio were ineffective at quantifying these morphological features of interest, reporting only on the increased elongation of the shape, but not other morphological changes. Because shapes with buds have increased perimeter, they are best quantified with circularity. Although concave regions are present in all of these shapes, the decrease in convexity is less pronounced in shapes 8–10 than in shapes with a greater number of protrusions (shapes 5–7) (**fig 1B**).

Collectively, these results suggest that circularity is best suited to quantify spheroids/organoids with isolated invasion tips or points, owing to its greater sensitivity to increases in perimeter relative to increases in area. Solidity and convexity, which detect deviations from the ideal convex contours, are less suited to describe such shapes. Finally, roundness and aspect ratio can detect bulk elongation of a shape but are agnostic to corners/points/invasion tips and, therefore, not a strong descriptor of the types of invasions formed by matrix-embedded spheroids/organoids (**fig 1B**). An image-resolution sensitivity analysis shows that measured shape descriptors remain consistent across different image sizes and pixel counts along the shape perimeter, validating their size independence and suitability for comparisons across length scales (**Supp fig 1**).

In research settings, invasive spheroids/organoids typically exhibit distinct invasion tips or fingers, which can be detected by segmenting regions of interest (ROIs, black **fig 1C**) from micrographs collected by different imaging modalities including bright-field, confocal, and optical coherence microscopy (**fig 1C**). Given the functional relevance of invasive protrusions, we designed additional digital phantoms with varying protrusion-to-diameter ratios to further probe the sensitivity of circularity, solidity, and convexity to detect structures consistent with small (tips) or larger invasions (fingers) (**fig 1D**). Indeed, all three shape descriptors decreased with an increased protrusion to diameter ratio, although the effect was greatest for circularity.

Next, we compared how well the different shape descriptors detected invasion in experimental images. To this end, we chose three sample spheroids that varied in morphology and invasion: round, non-invasive; elongated, non-invasive; and round, invasive (**fig 1E-F**). All spheroids were stained with phalloidin to visualize the F-actin cytoskeleton, imaged with confocal microscopy, 3D data processed into maximum projections, and segmented in FIJI into ROIs for shape factor analysis (**fig 1E-F**). Circularity, solidity, and convexity, all decreased for the round, invasive spheroid relative to its round, non-invasive and elongated, non-invasive counterparts (**fig 1F**). Similarly, circularity, solidity, and convexity decreased for the elongated, non-invasive vs. round, non-invasive spheroid, but was accompanied by a much more marked decrease in roundness relative to the round, invasive spheroid. Correspondingly, aspect ratio for the non-invasive, elongated vs. both non-invasive and invasive round spheroids increased significantly (**fig 1F**). Collectively, results from our comprehensive comparison of shape factors - using both digital phantoms and experimental images - suggest that circularity and convexity are effective at quantitatively differentiating non-invasive spheroids from invasive spheroids. In contrast, roundness can be used to assess sample elongation but is not a good metric for invasion. As aspect ratio is mathematically defined as the inverse of roundness, we have excluded it from our analysis from here on as it provides no additional distinctive shape descriptor value.

### Extending shape factor analysis to cancer organoids

Our analyses above suggested that different shape factors vary in their capacity to capture distinct morphological features, which not only affects the analysis of invasive spheroids, but is particularly pertinent when studying organoids. More specifically, organoids often exhibit concave regions associated with local folding/buckling or protrusions caused by non-symmetrical growth, which can be challenging to detect. To investigate the capability of each shape factor to assign quantitative morphological scores to organoids, we analyzed endometrial epithelial and oviductal tubal epithelial organoid cultures.

Endometrial cancer organoids were formed by isolating endometrial epithelial cells from the mouse uterus of *Trp53* and *Rb1* floxed mice, treating them with adenovirus-Cre, and culturing in Matrigel. Bright-field microscopy of non-fixed samples revealed that some organoids exhibited uniform circular shapes while others formed irregular shapes with protrusions, folds, or microlobules (**fig 2A**). Manual segmentation of four representative organoids of each category yielded ROIs for subsequent analysis. Interestingly, circularity was best able to differentiate irregularly shaped from uniformly shaped organoids as it detected the largest numerical differences between both categories (**fig 2B**). In contrast, solidity, convexity, and roundness were much less effective at identifying shape differences between conditions in this data set (**fig 2B**).

In addition to their gross morphologies, the size and structure of organoid lumens are also important as they can, for example, be indicative of malignancy. To test how well the different shape factors can detect cancer-related changes in organoid lumen morphology, carcinogenesis was induced in oviductal tubal epithelial cells via infection with adenovirus-Cre that inactivated *Trp53* and *Rb1* prior to organoid formation while control cells were infected with a blank adenovirus. Microscopic imaging of H&E-stained organoid sections revealed clusters of cells inside the lumen of cancer organoids but not controls (**fig 2C**). Inspired by prior studies quantifying lumen shape^29,30^, we manually segmented the internal cell-free lumens to generate ROIs for shape factor analysis. Due to the accumulation of cells, lumens of cancer organoids had concave structures, which predictably yielded lower values of convexity and solidity than the control organoids with convex lumens (**fig 2D**). Circularity was best able to quantify differences between the control and cancer organoid lumen shapes given the increase in concave vs. convex lumen perimeter (**fig 2D**). Of note, roundness did not change between the control and cancer organoids because the lengths of major and minor axes remained relatively unaffected by luminal cell growth. These quantitative results suggest that circularity may be the most universal ImageJ/FIJI shape factor to quantify changes in organoid morphology, but depending on the morphological features of interest (particularly folds and intra-luminal growths), solidity and convexity could also be of value. Integrating shape factor analysis into the analysis pipeline of organoids could help identify how organoid morphologies are determined by their genetic and lineage profiles^31^, potentially establishing a more easily accessible metric to characterize organoids in high-throughput settings.

### Quantifying collective invasion with radial length analysis

Although the results above suggest that certain basic shape descriptors are well-suited to characterize changes in spheroid invasion and organoid morphology, only circularity can be used relatively universally. Therefore, we sought to identify a more robust and specific metric. Based on the irregularities of their margins, radiologists categorize breast tumors into five main classes using the BI-RADS©^12^ system (**fig 3A)**. To describe these margin irregularities quantitatively ^21^, several approaches have been developed, including analysis of radial lengths described in 1993 by Kilday *et al.* ^25^. In this method, radial lengths are defined as the Euclidean distance from the center of the tumor to each boundary coordinate of the segmented shape. Variations of this radial length analysis have been used on multiple types of clinical breast or liver images^32–36^. To evaluate the suitability of this method to evaluate the morphology of *in vitro* cancer models, we have developed an analysis pipeline in FIJI and MATLAB that quantifies the standard deviation of radial lengths (SD_RL_) and the number of average radial length crossings (ARLC) in spheroids.

To illustrate the fundamentals underlying radial length analysis, we generated a fictional ROI of a star (**fig 3B**). In practice, ROIs are generated via automated or manual segmentation of the spheroid/organoid from images in FIJI and saved as the “.roi” file type. Subsequently, ROIs are imported into MATLAB^37^, the centroid of the shape is computationally determined, and radial lengths are found by calculating the Euclidean distance between the centroid and each point on the shape perimeter (**fig 3B**). The resulting radial length vector is plotted as a function of perimeter point index along with the computationally determined average radial length. The number of crossings or intersections between the two lines is determined and reported as the number of ARLC (**fig 3B**). This value corresponds geometrically to the number of undulations at the shape margin with respect to the centroid of the shape. The criteria for determining a crossing can be set and optimized depending on the noisiness of the data set. The two parameters that can be tuned are 1) the excursion length, which represents the number of consecutive perimeter points that must fall either above or below the average radial distance, and 2) the minimum crossing distance, which describes how far above/below the average the radial length vector points must be to count as a crossing (**fig 3B, supp fig 2**). Adjusting these parameters increases the signal to noise ratio and sets a minimum crossing threshold tuned to the scale of the protrusions for a specific experimental system or imaging resolution (**supp fig 2**). Finally, the SD_RL_ are calculated and reported as a metric indicating perimeter variations through either protrusions or an asymmetric/elongated shape. While ARLC are size independent, SD_RL_ are dependent on the resolution and/or scale of the image (**supp fig 3**) with default units of pixels unless converted back to length units as encoded by the original image.

**Figure 2:**
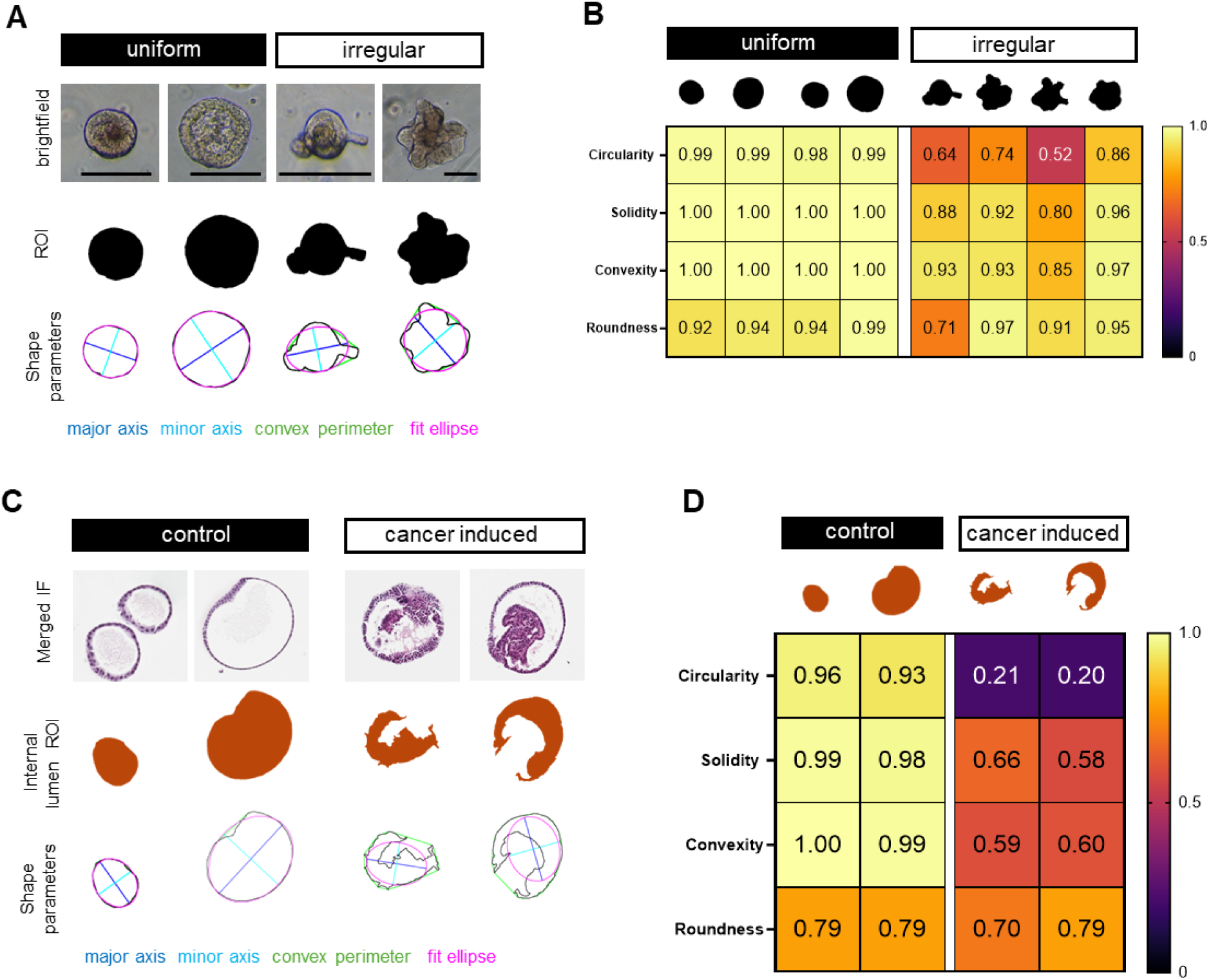
Shape factor analysis of organoids. **A**) Bright-field images of non-fixed, Matrigel-embedded endometrial epithelial organoids categorized as uniform or irregular and their corresponding manually segmented ROIs, as well as shape parameters for shape factor calculations. Scale bars = 150 µm. **B**) Heat map showing how FIJI-based shape factors vary between uniform and irregular organoids. **C**) H&E-stained cross-sections of control or cancer mouse oviductal tubal epithelial organoids. Internal lumens were manually segmented and shape parameters quantified to identify differences in luminal space morphology. All sections were scanned with the ScanScope CS2 (Leica Biosystems, Vista, CA) with the doublet inserted for 40x objective magnification) cropped images are displayed here for visualization. **D)** Heat map showing how FIJI-based shape factors of internal lumens vary between control and cancer-induced tubal epithelial organoids.

**Figure 3:**
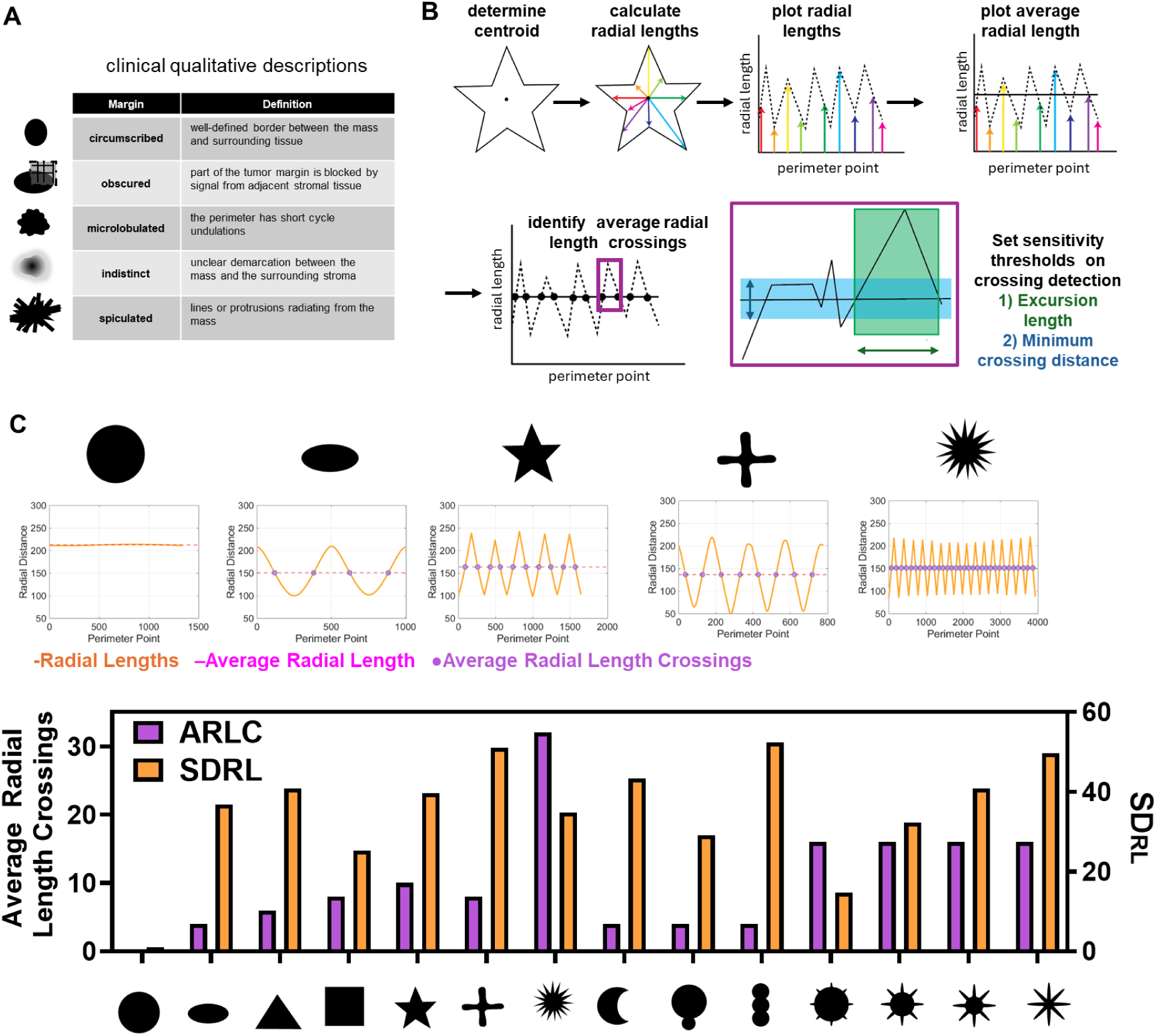
Evaluating radial length analysis as a quantitative shape descriptor on digital phantoms. **A)** Summary of clinical descriptors of breast tumor margins from the BI-RADS© system. **B)** Workflow of the computational methods used to determine variations in radial length and the number of average radial length crossings, including a visual depiction of the sensitivity parameters that can be adjusted for each data set. **C)** Application of the radial length analysis to digital phantoms and their corresponding radial length plot. The radial length vectors are displayed in orange with the average radial length shown in magenta and the average radial length crossings shown as purple points on each plot. Bar plot displays the number of average radial length crossings (ARLC) and the standard deviation of the radial lengths (SD_RL_).

To assess the capabilities of radial length analysis, we first applied the code to analyze the digital phantoms from figure 1 (**fig 3C, supp fig 3**). This analysis provided the following insights: 1) ARLC and SD_RL_ are independent metrics which combine to provide comprehensive quantification of shape geometries; 2) ARLC increase with membrane irregularities, but also with sharp corner geometries such as the square; 3) SD_RL_ increases with deviation from a circle either in higher aspect ratio or in protrusion length/number (**fig 3C**). ARLCs are not sensitive to differences in protrusion lengths for the tested geometries, but SD_RL_ increases with increasing protrusion length, indicating that both metrics should be combined when quantitatively classifying objects and thus, 3D culture models, using this approach (**fig 3C**). We found that for a given excursion length and minimum crossing distance, the number of ARLC was independent of image resolution while SD_RL_ increased proportionally to resolution in agreement with the size (in number of pixels) dependence of SD_RL_ (**supp fig 3B**).

### Spheroid ARLC analysis and adaptations for asymmetric invasion

After proof-of-concept testing with the digital phantoms, we next performed radial length analysis of the experimental spheroids shown in figure 1. We found that in invasive spheroids, both ARLC and SD_RL_ increased, but in elongated, non-invasive spheroids only SD_RL_ increased (**fig 4A-B**). These results were obtained using a minimum excursion length of 3 points and a minimum crossing distance of 5 pixels (**fig 4A-B**). Increasing both parameters causes a reduction in ARLC as the sensitivity of crossing detection is reduced, but the trends between conditions are preserved in all cases (**supp fig 4)**. Certain biological questions or organoid shapes may benefit from quantifying confocal cross-sections instead of maximum projections as used in **fig 4**. To demonstrate this potential application, we separately segmented ROIs from individual confocal images (10 µm slices, 50 µm between images shown sequentially in **supp fig 5A**). Results reveal small depth-dependent fluctuations in shape descriptors (**supp fig 5B**) with more prominent changes in ARLC at discrete depths vs. the maximum projection (**supp fig 5B**). Our pipeline can be readily used to quantify ARLC of whole spheroids or optical slices of radially invading spheroids.

**Figure 4:**
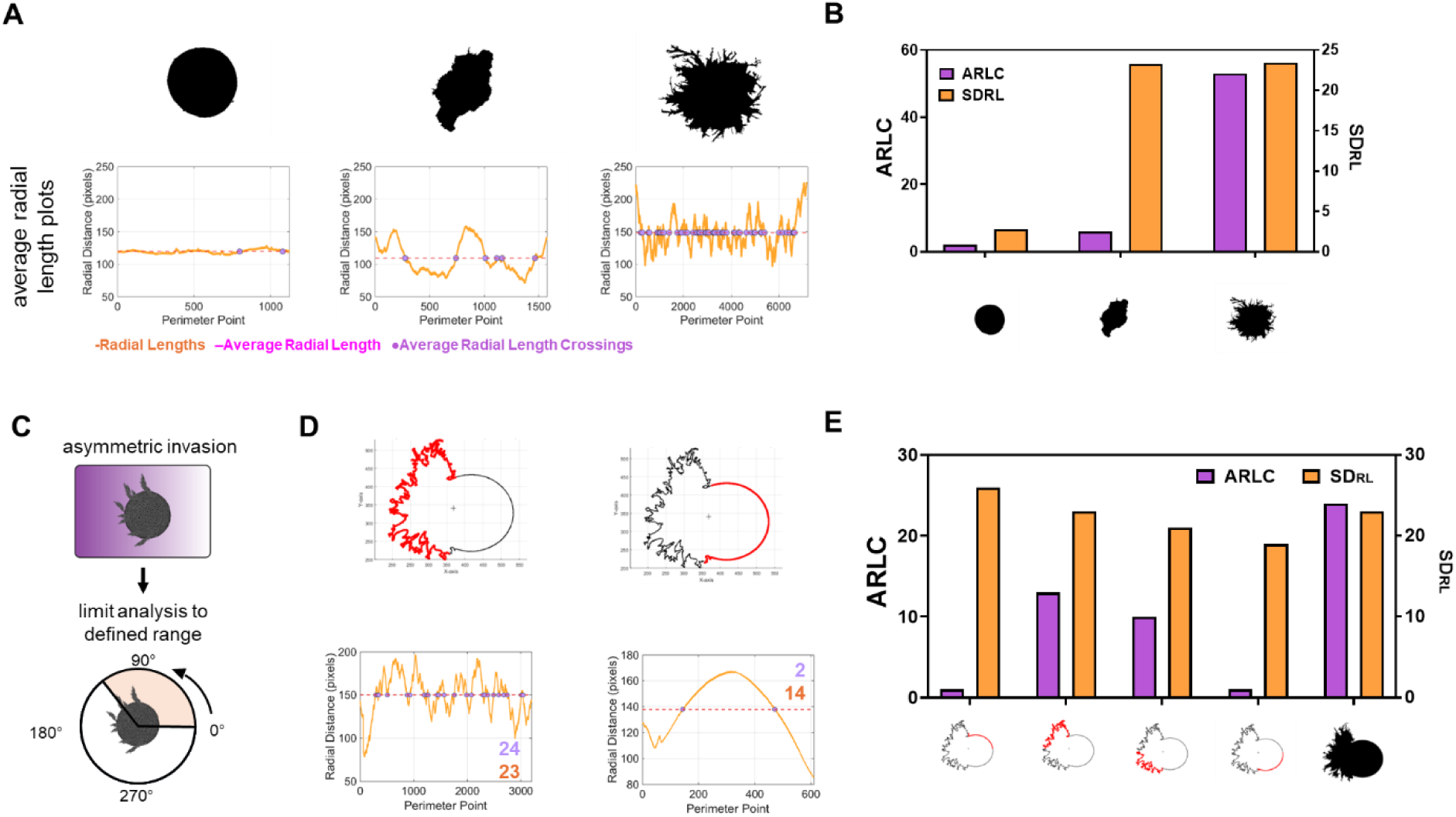
Applying radial length analysis to spheroids and implementing segment analysis for asymmetric invasion. **A)** Radial length analysis of sample spheroid ROIs demonstrates an ability to quantify border irregularities to separate non-invasive spheroids (left, middle) from invasive spheroids (right). The radial length vectors are displayed with orange with the average radial length shown in magenta and the average radial length crossings shown as purple points on each plot. **B)** Bar plot displays the number of average radial length crossings (ARLC, left axis) and the standard deviation of the radial lengths (SD_RL_, right axis,). **C)** Schematic of segmented radial length analysis where angles can be set to analyze a defined radial segment of the perimeter for analysis. **D)** Analysis of two segments of a digital phantom of asymmetric invasion where two halves (265°- 85°, left) and (85-265°, right). The segment of the spheroid to be analyzed is highlighted in red in the generated ROIs plots (top row), and the radial length plots are displayed below (bottom row) with ARLC annotated in purple and SD_RL_ annotated in orange text. **E)** Bar plot displaying the ARLC (purple, left axis) and SD_RL_ (orange, right axis) for 4 quadrants of the phantom plotted from left to right: 0-90°, 90-180°, 180-270°, and 270-360°and the whole ROI 0-360°.

**Figure 5:**
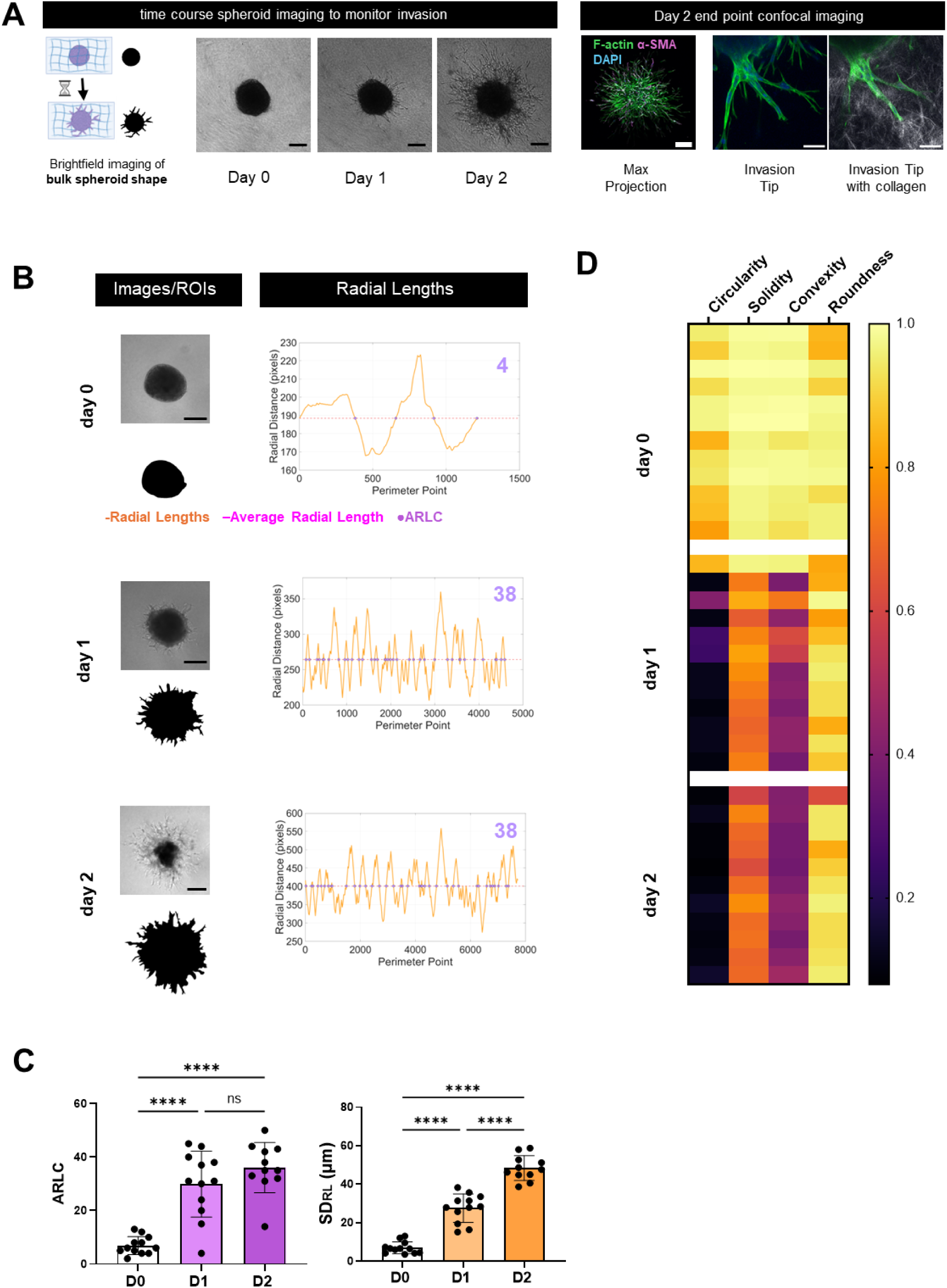
Longitudinal analysis of spheroid invasion with shape factor analysis. **A**) Schematic and representative images of spheroids prepared from adipose stromal cells and embedded into collagen type I hydrogels. Spheroids were imaged with bright-field microscopy every day to monitor invasion over time (left panel scale bar = 200 µm). Confocal images of fixed samples on day 2 reveal cellular level insights into collective cell invasion tips and migration through the surrounding hydrogel at the end point of the experiment. Scale bar = 200 µm max projection, = 50 µm in the invasion tip zoom view. **B**) Spheroid ROIs were isolated from day 0, day 1, and day 2 images to quantify changes in shape factors over time during the 48 hours of culture. Representative images and their corresponding ROIs and radial length plots are displayed. Crossing sensitivity set at minimum excursion length of 3 and minimum crossing distance of 5 pixels. **C**) Quantification of the ARLC and SD_RL_ show a significant increase when compared to day 0 in both parameters after both 1 and 2 days of culture. Statistical significance determined by unpaired t test with Welch’s correction (**** = p<0.0001, n=12). **D**) Heat map display of shape factors reveals spheroid circularity, solidity, and convexity decrease after 1 day of invasion while roundness remains constant. These trends are preserved into day 2. Schematic in A created in Biorender.com

Another aspect to consider is that asymmetric radial invasion (**fig 4C**) may occur in the presence of biochemical or physical microenvironmental gradients, driven by processes such as chemotaxis, durotaxis, or directed migration in response to matrix microarchitecture^38–41^. To optimize our pipeline to address this possibility, we adapted our code to analyze user defined radial segments or “pie slices” of the ROI (**fig 4C**). In this analysis, 0 degrees is set at the 3 o’clock position of the ROI and angles increase counterclockwise. We generated non-symmetrically invading digital phantoms (**supp fig 6**), analyzed 180-degree segments highlighted in red, and observed a higher number of ARLC in the irregular half (**fig 4D**, left) when compared to the half with smooth surface contours (**fig 4D**, right). This example also highlights a notable difference in the perimeter points required to define each half with over 3 times as many points required for the irregular left side of the phantom (**fig 4D**). Splitting the phantom into 90-degree quadrants reveals consistently high SD_RL_ across quadrants, but elevated ALRC values are limited to the quadrants containing irregular surfaces (**fig 4E**). In contrast, the whole phantom ROI (360 degrees perimeter, **fig 4E**) has both elevated ALRC and SD_RL_ which obscures the additional directional information that is gained from a segment analysis. Therefore, regions of spheroids/organoids should be isolated to investigate spatial variance in invasion patterns in observable cases of asymmetry to gain additional directional information.

**Figure 6:**
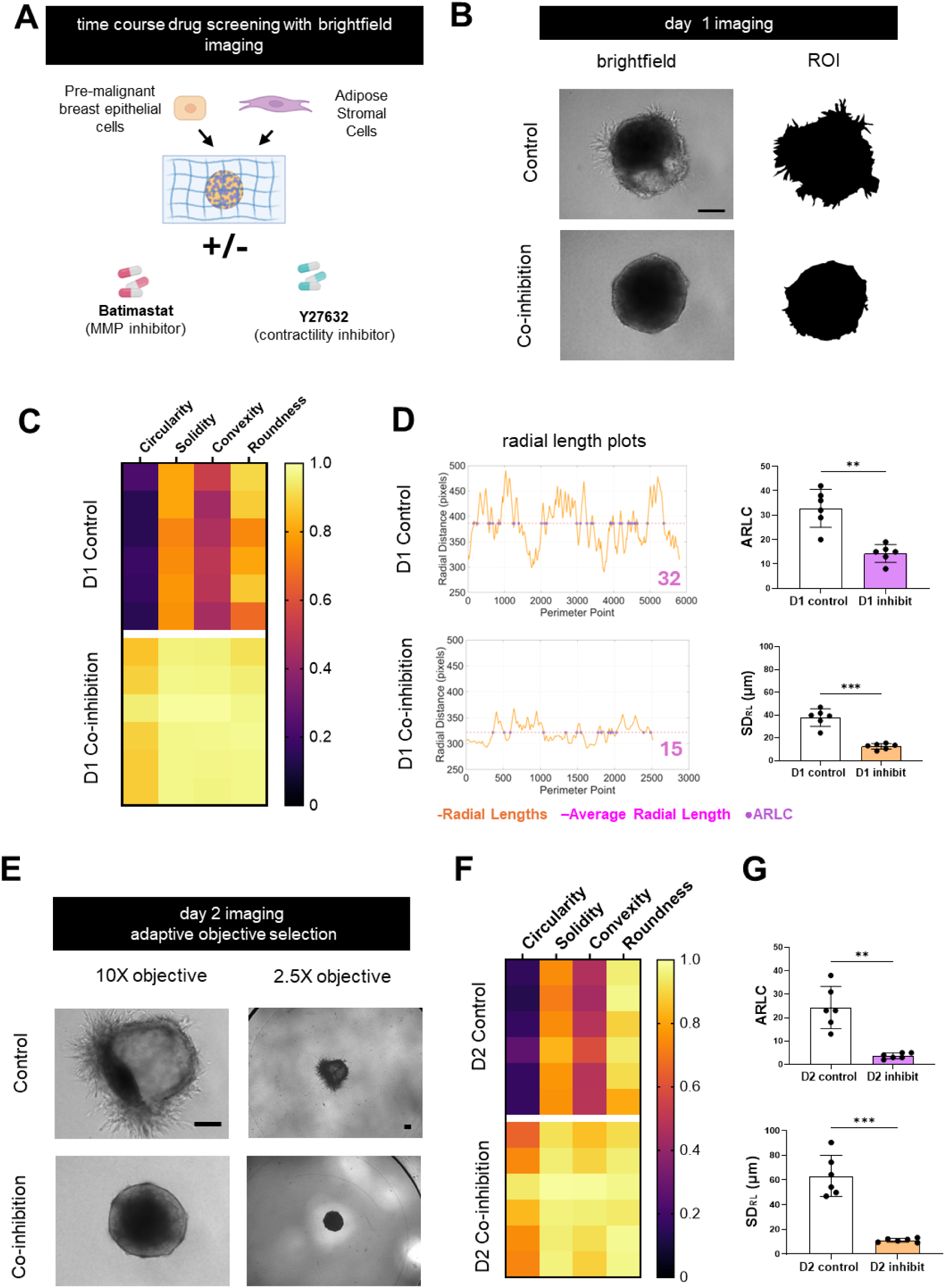
Radial length analysis of drug-treated spheroids identifies differences in invasion earlier and independently of objective magnification. **A)** Schematic of the experimental setup of collagen embedded co-culture spheroids of premalignant breast epithelial (MCF10AT1) and adipose stromal cells treated with a combination of Batimastat and Y27632 to co-inhibit matrix degradation and contractility. **B**) Representative images after 24 hours of inhibition and corresponding ROIs. **C**) Shape factor analysis demonstrates increased circularity, solidity, convexity, and roundness when spheroids were treated with inhibitors for 1 day. **D**) Average radial length crossings and the standard deviation of the radial lengths were significantly reduced with co-inhibition for 1 day. **E**) Images of day 2 spheroids taken with a 10X and 2.5X objective. Scale bars = 200µm. **F**) Shape descriptor analysis of day 2 spheroids with ROIs manually isolated from images taken with a 2.5X objective to ensure the whole spheroid was in the field of view. **G**) Radial length analysis of the day 2 spheroid ROIs from the 2.5X objective images. For D and G, n=6, * = p<0.05 as determined by an unpaired T-test with Welch’s correction. Schematic in A created in Biorender.com

### Live monitoring of spheroid invasion via shape factor analysis

Next, we performed shape factor analyses on a well characterized experimental system. To interrogate the invasive phenotypes of adipose stromal cells (ASC) and their role in breast cancer progression, we have previously embedded ASC spheroids into collagen hydrogels and monitored them through daily, low resolution, bright-field imaging (**Fig 5A, left**)^18^. At the end of the experiments, we also fixed and stained the samples for high resolution confocal or multiphoton imaging to better understand the relationship between ASC invasion and matrix remodeling (**Fig 5A, right**). However, the latter approach is costly, inherently low-throughput, time-intensive, and prohibits longitudinal imaging of shape changes. Instead, bright-field monitoring of live spheroids at set time points during the culture period may be able to capture dynamic morphological changes and allows for longitudinal and high-throughput analysis.

To investigate if either ARLC analysis or FIJI shape descriptors could detect invasion as early as one day following spheroid embedding, we re-analyzed our previously published data set for which we had examined invasion by simply measuring changes in spheroid area. Radial length analysis of manually generated ROIs from 12 spheroids demonstrates that ARLC and SD_RL_ of spheroids increased after 1 day in culture, enabling a more precise discrimination of emerging invasive phenotypes than our previous analysis **(fig 5B-C**). On day 2, ARLC were not statistically different from day 1, but we observed an increase in the SD_RL_, which could explain the change in spheroid area between days 1 and 2 that we detected previously^18^. Similarly, basic shape factor analysis reveals striking differences emerging as early as day 1, with decreases observed across all metrics except for roundness (**fig 5D**). Circularity and convexity mirror the trends observed in ARLC with large differences between day 0 and 1 and minimal differences between day 1 and 2 (**fig 5D**). In summary, ARLC, circularity, and convexity all detect day 1 differences, highlighting the utility of shape factor analysis for longitudinal analysis of invasion in non-fixed samples via bright-field microscopy. However, ARLC analysis is the only metric that specifically detects radial protrusions associated with the observed collective invasion phenotype.

### Shape factors analysis supports time-resolved drug screening studies

Lastly, we evaluated if shape factor analysis could be used to detect how a model drug regimen suppresses spheroid invasion. In our previous work, we treated co-culture spheroids of pre-malignant breast cancer cells and ASCs with Batimastat, a matrix metalloproteinase inhibitor, and Y27632, a contractility inhibitor^18^ (**fig 6A**). Endpoint imaging showed that ASCs promoted invasion of pre-malignant breast cells, but treatment with the drug combination prevented invasion^18^. These results were obtained by evaluating changes in spheroid area using images obtained by bright-field microscopy, but the detected differences were relatively small. Revaluating the data set by creating ROI masks generated at day 1 visually illustrates the striking reduction in perimeter irregularities in inhibitor-treated spheroids (**fig 6B**). All tested shape descriptors quantify this difference between control and co-inhibition; however, circularity and convexity quantitatively discriminate the conditions the most (**fig 6C**). These results can be interpreted as drug treatment preventing the formation of collective invasion tips that increase the perimeter (decreasing circularity) and create concave regions (decreasing convexity and solidity). Additionally, uninhibited asymmetric invasion can lead to spheroid elongation (decreased roundness). As previously determined, these metrics are not specific to invasion and are highly impacted by confounding variables and morphological heterogeneity. The invasion specific radial length analysis revealed statistically significant reductions in ARLC and SD_RL_ with drug treatment at day 1 (**fig 6D**). The advantage of the dimensionless nature of shape factor analysis also allows for comparisons across time points and with imaging setup variation in objectives, resolutions, and instruments (**fig 6E**). This is true for all metrics used here except for SD_RL_ which does have units that correspond to physical dimensions (in this case µm). This feature is particularly important when invasion area of spheroids in one condition or time point is much more pronounced than in others and thus requires switching of objectives between conditions. For example, control spheroids on day 2 had to be imaged at lower magnification as compared to day 0 and 1 due to their extensive invasion (**fig 6E**). Analyzing the low magnification day 2 samples show the same trends as the higher magnification and resolution images on day 1. The dimensionless nature of ARLC allows the day 1 values (**fig 6C-D**) to be directly compared to the day 2 values in (**fig 6F-G**) despite the differences in image acquisition parameters. Overall, shape factor analysis identified drug-dependent differences in invasion earlier than simply analyzing spheroid area, opening the possibility of additional high throughput data collection and analysis in future studies.

## Discussion and Conclusion

Linking tissue shape and structure to biological function remains a challenge across both clinical practice and bench science. Radiologists undergo extensive training to discriminate and score subtle shape differences in clinical imaging. Researchers studying spheroids and organoids would benefit from comparable expertise but currently lack equivalent tools and standardized frameworks to reliably interpret morphology. As organoid and spheroid models become central to disease modeling and pre-clinical screening^9^, scalable, quantitative methods to interpret complex morphology are required. Inspired by computer assisted breast imaging technologies, we sought to apply dimensionless shape factor analysis to the morphological quantification of 3D *in vitro* model systems through a simple and user-friendly workflow in FIJI and MATLAB. We first developed a series of digital phantoms to serve as a visual guide to demonstrate which shape factors detect specific target morphologies. Next, we provided multiple experimental examples to assist users in selecting which metrics may be best suited for future interpretation of their own biological data sets.

FIJI and CellProfiler^42,43^ include a subset of user-friendly dimensionless shape factors that allow comparisons across length scales, imaging modalities, and variation in experimental conditions. Shape factors have successfully been used to classify single cell morphology and serve as a predictive marker of a tumorigenic cell phenotype^44,45^. Because of the simplicity and broad relevance, FIJI shape descriptors can be readily applied beyond the subfield of cancer research. For example, in developmental biology, shape classification of brain^46^, intestinal^47,48^, or kidney^30^ organoids can provide critical information on morphogenic processes^49^. The computational shape classifications proposed here could additionally be used in organoid experiments for disease modeling (kidney^50^, pancreas^51^, heart^52^), reproductive health (ovarian tissue^53^), and toxicity screening^9^ to generate hypotheses linking morphology to biological phenotype.

In the context of cancer, we have focused on multicellular spheroid invasion assays; however, shape factor analysis would be useful much earlier in the spheroid experimental workflow, e.g. to validate which fabrication method is best suited to generate uniformly shaped spheroids. Indeed, prior work has demonstrated variations in spheroid structure imparted by the fabrication method can impact gene expression and drug sensitivity^54^. While we have focused here on applications in *in vitro* models, shape factor analysis could also prove useful in *in silico* experimental systems^55^. Our matrix invasion analyses show that convexity and circularity can quantitatively capture shape features linked to collective invasion by identifying increased perimeter to area ratios and concave regions that would result from invasive fingers. However, they are confounded by features such as a single large protrusion and may fail to detect small-scale roughness from invasion tips. Solidity, convexity, and circularity all decrease with non-invasive geometries such as elongation, an indented concave region, or a budding structure, thus limiting the interpretation of results when comparing to decreased values also observed in invasion. Despite the colloquial use of round to describe a shape with no sharp corners, it is mathematically defined as the inverse of aspect ratio, capturing only elongation and making it unsuitable for most spheroid and organoid analysis applications

We created the MATLAB radial length analysis to address the FIJI shape descriptors’ poor specificity for margin irregularities. Our custom-built code includes tunable inputs to define the size and scale of perimeter variations that would be defined as invasive tips or fingers in a specific biological context. When applied to a data set of collectively invading spheroids, the number of ARLC successfully differentiated spheroids based on invasiveness and roughness of the surface contours (**fig 4-6**). ARLC is best suited for identifying symmetric radial invasion or unidirectional spheroid invasion (segmented approach) in collectively invading spheroids. Spheroids with predominantly single cell invasion would benefit from alternative segmentation and analysis methods to independently evaluate single cells vs. the bulk spheroid core^56,57^. Similarly, 3D cell culture systems designed for predominantly non-spheroidal unidirectional single or collective cellular invasion ^58–61^ are not suited for bulk tissue-level shape factor analysis; however, applying shape factor analysis to individual cells or individual nuclei could uncover phenotypic heterogeneity among invading cells.

The simplicity of the ARLC analysis imposes several limitations that will require further method development and algorithm refinement. One confounding factor is ROI elongation whereby ARLC will decrease and SD_RL_ will increase in an elongated shape independent of the presence/absence of surface irregularities. Additionally, the current ARLC calculation has sensitivity limits on small perimeter undulations with respect to the overall diameter and limited interpretation of complex features such as bifurcated invasion tips. The radial segmentation process assigns perimeter points to a radial pie slice only based on the angle of the point with respect to the centroid. Therefore, this method cannot account for parts of invasion fingers that distally curl to extend past the defined radial slice, resulting in a loss of data. The ARLC code uses a single value of the centroid and ARL calculated from the whole ROI that is subsequently applied to each radial slice. Computing a new average radial length and a new centroid for each segment would serve to more accurately quantify the variations present in that segment independent of the rest of the spheroid, where for example, a very long average radial length may skew results for a segment that is all closer to the centroid than the average radial length. An alternative method to overcome this is to not calculate the radial length as the distance from the perimeter point to the centroid, but instead, calculate the radial deviation from a low-order contour approximation, using elliptical fourier decomposition which follows the bulk shape of the spheroid to emphasize protrusions across the full spheroid surface^62^. Similarly, fit ellipse metrics such as lobulation count^63^ have been used in hepatic imaging studies and should be tested in spheroids/organoids. The quantifications presented here become even more powerful when combined with automated organoid/spheroid segmentation assisted by deep learning or convolutional networks^64^. Examples of preexisting segmentation and analysis tools include ilastik^65^, INSIDIA^66^, SpheroScan^67^, and SpheroidJ^68^. For a summary of previously developed spheroid specific analysis and outputs please see the table compiled by Moriconi *et. al*.^66^. Applications with high throughput systems will also require quality control/outlier removal with tools such as SpheroidAnalyseR^69^. Using these existing spheroid/organoid segmentation tools first will automate the generation of ROIs that can then be analyzed with the shape factor tools presented here, ultimately increasing the analytical throughput for future applications. Combining radial length and shape factor analysis with tools to analyze cellular and subcellular changes^18,21,70–73^ will advance overall understanding of the mechanisms involved in invasion. Moreover, such studies could identify morphological biomarkers that enable imaging of invasion at low resolution and magnification at multiple time points in a large number of samples as, for example, necessary in platforms with robotic liquid handling and spheroid culture^74^ for precision medicine applications.

In summary, we have analyzed the capabilities and limitations of FIJI shape descriptors and compared them with radial length analysis to detect specific morphological features of shapes. Through the analysis of digital phantoms, organoids, and invasive spheroids, we have demonstrated the potential biological applications of morphology analysis and outlined key criteria for selecting shape factors. Developing automated, interpretable shape analysis pipelines would bridge the expertise gap between clinical image interpretation and laboratory practice, improving reproducibility, accelerating translational discovery, and allowing morphology-based phenotyping at the throughput required for large scale applications such as drug testing.

## Methods

### Creation of digital phantoms

Phantoms were hand drawn as regions of interest (ROIs) in FIJI using the oval, polygon, or freehand selection tools on a new 512×512 8-bit image^24^. ROIs were interpolated with 1 pixel interval to set the number of perimeter points. For resolution analysis, generated ROIs were scaled by 0.5 or 2 to proportionally fit 256×256 and 1024×1024 8-bit images respectively and interpolated at their new size. Asymmetric phantoms featured in supplemental figure 6 were generated through a combination of hand drawn original shapes and modification of experimental samples. Shapes were imported into FIJI, ROIs generated via the analyze particle tool, and interpolated with a 1-pixel interval.

### Primary mouse endometrial and oviductal tubal epithelial organoid culture

Primary cells were harvested from the mouse uterus or oviduct of *Trp53* and *Rb1* floxed mice^75,76^. Endometrial organoids were generated from single endometrial epithelial cells cultured in Matrigel and treated with adenovirus-CRE at the time of isolation to induce a cancer phenotype. Non-fixed organoids were imaged on day 7 in the Matrigel culture. A fraction of the organoids was treated with 4 days of DBZ; however, treatment vs. control conditions are not differentiated for the analysis and application in this shape factor study. Tubal epithelial organoids were generated via culture in Matrigel. For these organoids, control cultures were treated with an adenovirus blank while cancer induced organoids were infected with adenovirus-CRE that inactivated *Trp53* and *Rb1* in single cells on day 0 of culture. After 14 days of culture, organoids were isolated from Matrigel and resuspended in 4% paraformaldehyde for 1 hour. Fixed organoids were washed with 1x PBS 3 times prior to resuspending in HistoGel (Epredia, HG4000012) at 65 °C. HistoGel suspensions of organoids were set after 10 minutes, where they were then subjected to standard tissue processing and paraffin embedding^76^. 4µm paraffin sections of organoids were H&E stained for imaging.

### Collagen embedded adipose stromal spheroids, and adipose stromal-breast cancer co-culture spheroids

Murine adipose stromal cells were isolated from the mammary fat pads of wild type mice as previously described^18^. Human stromal cells were Adipose Derived Stem Cells (Lonza) and human breast cancer cells were the MCF10AT1 cell line or HAS3 overexpressing MCF10A cells^59^. Cells were expanded in their corresponding media^18,59^ and formed into spheroids of 5,000-10,000 cells per spheroids via aggregation on agarose coated plates on an orbital shaker for 24 hours or via the hanging drop method using media supplemented with 25cp Methylcellulose and type 1 collagen and cultured for 24 hours. Spheroids were embedded into 6mg/mL collagen type 1 hydrogels and cultured for 2-3 days with bright-field imaging every 24 hours. Spheroids were fixed in 4% PFA and stained with Alexa fluor 488 labeled phalloidin and DAPI before being imaged on a Zeiss 710 or 880 confocal microscope.

### Automated generation of spheroid regions of interest from fluorescence images

Confocal maximum projections of F-actin-stained spheroids were thresholded using the triangle thresholding algorithm in FIJI to create a binary image. The analyze particles tool was used to detect and create spheroid ROIs from these binary images, which were subsequently saved in the ROI manager. Spheroid images were acquired in 512×512 pixel images and ROIs interpolated to a 1-pixel interval. Detached single cells or clusters of cells were excluded from the ROIs to focus on collective invasion from the spheroid.

### Manual generation of spheroid/organoid regions of interest from bright-field images

Both bright-field images of spheroids/organoids and H&E stained cross sections of organoids were manually traced in FIJI to generated ROIs from the source images via the polygon selection tool. ROIs were interpolated with a 1-pixel interval.

### FIJI shape descriptors

FIJI shape descriptors were added to the acquired measurement parameters by checking the shape descriptors box in the set measurements menu of the software. ROI measurements yielded results for aspect ratio, roundness, circularity, and solidity – all of which are dimensionless units. Details of shape descriptor calculations can be found in the FIJI user guide^27^.

### Convexity quantification via a custom-built FIJI macro

Convexity is not a parameter conventionally included in the shape descriptor measurements of FIJI and therefore had to be generated using a FIJI macro. Briefly, the Convex Hull function was run to generate the ROI of the convex hull of the ROI. The convex hull is added to the ROI manager, measured to obtain its perimeter, and divided by the perimeter of the original ROI. Full code is available in the supplementary material.

### MATLAB script for radial length analysis

To determine the standard deviation of radial lengths (SD_RL_) and the number of average radial length crossings (ARLC) from ROIs generated from phantoms or experimental images, a MATLAB method was developed. Briefly, FIJI-generated ROIs were first imported into MATLAB using an adapted version of the ReadImageJROI plugin^37^ originally built by Dylan Muir (https://github.com/BrittanySchutrum/ReadImageJROI/blob/master/ReadImageJROI.m). Next, the centroid coordinates of the shape were obtained from the ROI by isolating the data from the adapted ROIs2Regions function^37^ (https://github.com/BrittanySchutrum/ReadImageJROI/blob/master/ROIs2Regions.m). The centroid and all perimeter points of the ROIs were plotted on a XY plot for visualization and conformation purposes. To determine the radial length from each perimeter point (i) to the centroid, the mean squared distance was determined as follows:

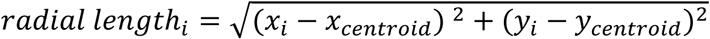

where x_i_ is the x coordinate of perimeter point i, y_i_ is the y coordinate of perimeter point i, and x_centroid_ and y_centroid_ are the x and y coordinate of the centroid, respectively. All the radial lengths were collected into a vector of the length of the number of perimeter points. The SD_RL_ was found from the standard deviation of this radial length vector. The average radial length – computed by calculating the average of the original radial length vector-was plotted as a horizonal line. The number of ARLC is computed by looking at pairs of data points - radial length_i_ and radial length_i+1_. A crossing is an instance where the radial length of the i point and i+1 point span the average, so in the case that 1) [radial length_i_ > average radial length & radial length_i+1_ < average radial length] or 2) [radial length_i_ <average radial length & radial length_i+1_ > average radial length]. Detection of crossings can be restricted by 1) a minimum number of points on each side of the average (defined as the excursion length) and 2) a minimum distance that the radial length must cross the average radial length by (defined as the minimum crossing distance). The developed code allows restricting crossing detection by excursion length and minimum crossing distance to reduce noise and fit the pixel length scale of the individual data set. Full MATLAB code is available in the supplemental material. Additional information and instructions are provided in the README file in the GitHub repository https://github.com/BrittanySchutrum/AverageRadialLengthCrossings-Spheroids.

### Data presentation

Heat maps and bar charts were generated in GraphPad Prism. All line plots generated in MATLAB. Schematics were drawn using Biorender, Adobe Illustrator, and PowerPoint.

### Statistical analysis

All statistical analysis was done in GraphPad Prism with the details of each statistical test provided in the corresponding figure captions.

## Supplementary Materials

Supplementary Materials include supplemental figures 1-6.

Figure list:

1. Shape descriptor image resolution sensitivity analysis of the basic shape digital phantoms
2. Sample data to illustrate how altering the parameters of the average radial length crossing quantification (excursion length and minimum distance) changes the detection of crossings
3. Digital phantoms evaluated by shape descriptors and radial length analysis
4. The impact of detection parameters and resolution of the sample spheroids on ARLC and SD_RL_
5. Depth dependent shape factor analysis of optical slices of a spheroid acquired via confocal imaging
6. Analysis of asymmetric digital phantoms with simulated directed invasion to the left

## Acknowledgements

The authors thank Dr. Rebecca Williams for her imaging expertise, review of this manuscript, and feedback throughout the process of developing the analysis used in this manuscript. We acknowledge financial support from NCI R01CA259195, R01CA276392, R01CA297166, and the Breast Cancer Alliance. Confocal imaging was conducted at the Cornell Biotechnology Resource Center Imaging Facility with the following funding: NIH S10RR025502 for the Zeiss LSM 710 confocal microscope and NIH S10OD018516 for the Zeiss LSM880 confocal microscope.

## Conflict of Interest

The authors have no conflicts to disclose.

## Ethics Approval

Animal studies were approved by the Cornell University Institutional Laboratory Animal Use and Care Committee protocol number 2000-0116.

## Author Contributions

**Brittany E. Schutrum**: Conceptualization, Formal Analysis, Methodology, Investigation, Software, Validation, Visualization, Writing/Original Draft Preparation **Jenny Deng**: Conceptualization, Methodology, Software, Validation, Writing/Review & Editing **Ju Hee Kim**: Formal Analysis, Validation, Visualization, Investigation **Amalie Gao**: Methodology, Software, Validation, Data Curation **Emily Hur**: Methodology, Conceptualization, Software, **Jack C. Crowley:** Methodology, Software **Lu Ling**: Investigation, Resources **Matalin G. Pirtz:** Investigation, Resources **Coulter Q. Ralston**: Investigation, Resources **Alexander Y. Nikitin**: Resources **Claudia Fischbach**: Conceptualization, Funding Acquisition, Methodology, Supervision, Writing- original draft, Writing – review & editing,

## Data Availability

The data that supports the findings of this study are available from the corresponding author upon reasonable request

## Code Availability

All code available on Github:

FIJI Macros https://github.com/BrittanySchutrum/FIJI-Spheroid-Morphological-Signatures DOI: https://doi.org/10.5281/zenodo.18881209

MATLAB Script https://github.com/BrittanySchutrum/AverageRadialLengthCrossings-Spheroids DOI: https://doi.org/10.5281/zenodo.18880960

## Notes

### Competing Interest Statement

The authors have declared no competing interest.

## References

1. Tan, M. L., Ling, L. & Fischbach, C. Engineering strategies to capture the biological and biophysical tumor microenvironment in vitro. Adv. Drug Deliv. Rev. 176, 113852 (2021).

2. Nunes, A. S., Barros, A. S., Costa, E. C., Moreira, A. F. & Correia, I. J. 3D tumor spheroids as in vitro models to mimic in vivo human solid tumors resistance to therapeutic drugs. Biotechnol. Bioeng. 116, 206–226 (2019).

3. Beeghly, G. F., Amofa, K. Y., Fischbach, C. & Kumar, S. Regulation of Tumor Invasion by the Physical Microenvironment: Lessons from Breast and Brain Cancer. 10.1146/annurev-bioeng-110220-115419 24, 29–59 (2022).

4. Berens, E. B., Holy, J. M., Riegel, A. T. & Wellstein, A. A cancer cell spheroid assay to assess invasion in a 3D setting. Journal of Visualized Experiments 2015, (2015).

5. Vinci, M., Box, C. & Eccles, S. A. Three-Dimensional (3D) Tumor Spheroid Invasion Assay. J. Vis. Exp. 2015, 52686 (2015).

6. Kunz-Schughart, L. A., Freyer, J. P., Hofstaedter, F. & Ebner, R. The use of 3-D cultures for high-throughput screening: The multicellular spheroid model. J. Biomol. Screen. 9, 273–285 (2004).

7. Li, Y. & Kumacheva, E. Hydrogel microenvironments for cancer spheroid growth and drug screening. Sci. Adv. 4, (2018).

8. Bregenzer, M. E., Mehta, P., Burkhard, K. M. & Mehta, G. Cracking the code: predicting tumor microenvironment enabled chemoresistance with machine learning in the human tumoroid models. npj Biomedical Innovations 2025 2:1 2, 28- (2025).

9. Wang, D., Villenave, R., Stokar-Regenscheit, N. & Clevers, H. Human organoids as 3D in vitro platforms for drug discovery: opportunities and challenges. Nature Reviews Drug Discovery 2025 1–23 (2025) doi:10.1038/s41573-025-01317-y.

10. Sung, H. et al. Global Cancer Statistics 2020: GLOBOCAN Estimates of Incidence and Mortality Worldwide for 36 Cancers in 185 Countries. CA Cancer J. Clin. 71, 209–249 (2021).

11. Harbeck, N. et al. Breast cancer. Nature Reviews Disease Primers 2019 5:1 5, 1– 31 (2019).

12. D’Orsi, C., Sickles, E., Mendelson, E., Morris EA & et al. ACR Bi-RADS© Atlas, Breast Imaging Reporting and Data System. (American College of Radiology, Reston, VA, 2013).

13. Surveillance Research Program, N. C. I. SEER*Explorer: An interactive website for SEER cancer statistics [Internet]. SEER Incidence Data, November *2022 Submission (1975-2020)* (2023).

14. Oza, P., Sharma, P., Patel, S. & Bruno, A. A Bottom-Up Review of Image Analysis Methods for Suspicious Region Detection in Mammograms. J. Imaging 7, (2021).

15. Masud, R., Al-Rei, M. & Lokker, C. Computer-Aided Detection for Breast Cancer Screening in Clinical Settings: Scoping Review. JMIR Med Inform 2019;7(3):e12660 https://medinform.jmir.org/2019/3/e12660 7, e12660 (2019).

16. Vong, C. K., Wang, A., Dragunow, M., Park, T. I. H. & Shim, V. Quantification of tumorsphere migration with a physics-based machine learning method. Cytometry Part A 103, 518–527 (2023).

17. Vaidyanathan, K. et al. A machine learning pipeline revealing heterogeneous responses to drug perturbations on vascular smooth muscle cell spheroid morphology and formation. Scientific Reports 2021 11:1 11, 1–15 (2021).

18. Ling, L. et al. Obesity-Associated Adipose Stromal Cells Promote Breast Cancer Invasion through Direct Cell Contact and ECM Remodeling. Adv. Funct. Mater. 30, 1910650 (2020).

19. Puls, T. J. et al. Development of a Novel 3D Tumor-tissue Invasion Model for High-throughput, High-content Phenotypic Drug Screening. Scientific Reports 2018 8:1 8, 1–14 (2018).

20. Perrin, L., Belova, E., Bayarmagnai, B., Tüzel, E. & Gligorijevic, B. Invadopodia enable cooperative invasion and metastasis of breast cancer cells. Communications Biology 2022 5:1 5, 1–14 (2022).

21. Ilina, O. et al. Cell–cell adhesion and 3D matrix confinement determine jamming transitions in breast cancer invasion. Nature Cell Biology 2020 22:9 22, 1103–1115 (2020).

22. Poddar, S. et al. Metadherin with Stromal-Immune Cues Drives CD36-Dependent Lipid Reprogramming and Metastasis in Triple-Negative Breast Cancer: Insights from a Hetero-Spheroid Model. Adv. Healthc. Mater. e04890 (2026) doi:10.1002/ADHM.202504890.

23. Elosegui-Artola, A. et al. Matrix viscoelasticity controls spatiotemporal tissue organization. Nature Materials 2022 22:1 22, 117–127 (2022).

24. Schindelin, J. et al. Fiji - an Open Source platform for biological image analysis. Nat. Methods 9, 10.1038/nmeth.2019 (2012).

25. Kilday, J., Palmieri, F. & Fox, M. D. Classifying mammographic lesions using computerized image analysis. IEEE Trans. Med. Imaging 12, 664–669 (1993).

26. Russ, J. & Neal, B. The Image Processing Handbook. (CRC Press, Taylor & Francis Group, Boca Raton, 2016).

27. Analyze Menu. https://imagej.net/ij/docs/menus/analyze.html.

28. Kopanja, L., Lončar, B., Žunić, D. & Tadić, M. Nanoparticle shapes: Quantification by elongation, convexity and circularity measures. Journal of Electrical Engineering 70, 44–50 (2019).

29. Indana, D. et al. Lumen expansion is initially driven by apical actin polymerization followed by osmotic pressure in a human epiblast model. Cell Stem Cell 31, 640–656.e8 (2024).

30. Beck, L. E. et al. Systematically quantifying morphological features reveals constraints on organoid phenotypes. Cell Syst. 13, 547–560.e3 (2022).

31. Phuong, D. J. et al. Combinatorial organoid mutagenesis screen reveals gene constellations driving malignant transformation, pathology and chemosensitivity in high-grade serous ovarian carcinoma. bioRxiv 2025.03.10.642422 (2025) doi:10.1101/2025.03.10.642422.

32. Nie, K. et al. Quantitative Analysis of Lesion Morphology and Texture Features for Diagnostic Prediction in Breast MRI. Acad. Radiol. 15, 1513–1525 (2008).

33. Sahiner, B., Chan, H.-P., Petrick, N., Helvie, M. A. & Hadjiiski, L. M. Improvement of mammographic mass characterization using spiculation measures and morphological features. https://doi.org/10.1118/1.1381548 (2001) doi:10.1118/1.1381548.

34. Tsui, P. H. et al. Classification of benign and malignant breast tumors by 2-D analysis based on contour description and scatterer characterization. IEEE Trans. Med. Imaging 29, 513–522 (2010).

35. Islam, W., Danala, G., Pham, H. & Zheng, B. Improving the performance of computer-aided classification of breast lesions using a new feature fusion method. 10.1117/12.2611841 12033, 98–105 (2022).

36. Xu, J., Faruque, J., Beaulieu, C. F., Rubin, D. & Napel, S. A Comprehensive Descriptor of Shape: Method and Application to Content-Based Retrieval of Similar Appearing Lesions in Medical Images. J. Digit. Imaging 25, 121 (2011).

37. Muir, D. R. & Kampa, B. M. FocusStack and StimServer: a new open source MATLAB toolchain for visual stimulation and analysis of two-photon calcium neuronal imaging data. Front. Neuroinform. 8, 107360 (2015).

38. Sharma, S. et al. Mechanical cues guide the formation and patterning of 3D spheroids in fibrous environments. PNAS Nexus 4, 263 (2025).

39. Ayuso, J. M. et al. Study of the Chemotactic Response of Multicellular Spheroids in a Microfluidic Device. PLoS One 10, e0139515 (2015).

40. Ascione, F. et al. Gradient-induced instability in tumour spheroids unveils the impact of microenvironmental nutrient changes. Scientific Reports | 14, 20837 (123AD).

41. Geiger, F., Schnitzler, L. G., Brugger, M. S., Westerhausen, C. & Engelke, H. Directed invasion of cancer cell spheroids inside 3D collagen matrices oriented by microfluidic flow in experiment and simulation. PLoS One 17, e0264571 (2022).

42. Broad Institute. Welcome to CellProfiler’s documentation!

43. Stirling, D. R. et al. CellProfiler 4: improvements in speed, utility and usability. BMC Bioinformatics 22, 1–11 (2021).

44. Baskaran, J. P. et al. Cell shape, and not 2D migration, predicts extracellular matrix-driven 3D cell invasion in breast cancer. APL Bioeng. 4, (2020).

45. Conner, S. J. et al. Cell morphology best predicts tumorigenicity and metastasis in vivo across multiple TNBC cell lines of different metastatic potential. Breast Cancer Research 26, 43- (2024).

46. Boerstler, T. et al. Deciphering brain organoid heterogeneity by identifying key quality determinants. Communications Biology 2025 8:1 8, 1–11 (2025).

47. Montes-Olivas, S. et al. In-silico and in-vitro morphometric analysis of intestinal organoids. PLoS Comput. Biol. 19, e1011386 (2023).

48. Yavitt, F. M. et al. In situ modulation of intestinal organoid epithelial curvature through photoinduced viscoelasticity directs crypt morphogenesis. Sci. Adv. 9, eadd5668 (2023).

49. Gritti, N. et al. MOrgAna: accessible quantitative analysis of organoids with machine learning. Development 148, (2021).

50. Freedman, B. S. et al. Modelling kidney disease with CRISPR-mutant kidney organoids derived from human pluripotent epiblast spheroids. Nat. Commun. 6, 8715 (2015).

51. Below, C. R. et al. A microenvironment-inspired synthetic three-dimensional model for pancreatic ductal adenocarcinoma organoids. Nat. Mater. 21, 110–119 (2022).

52. Zhao, Y. et al. Geometrically controlled cardiac microtissues promote vascularization and reduce inflammation in vitro and in vivo. Cell Biomaterials 1, 100075 (2025).

53. Nason-Tomaszewski, C. E., Thomas, E. E., Matera, D. L., Baker, B. M. & Shikanov, A. Extracellular matrix-templating fibrous hydrogels promote ovarian tissue remodeling and oocyte growth. Bioact. Mater. 32, 292–303 (2024).

54. Gencoglu, M. F. et al. Comparative Study of Multicellular Tumor Spheroid Formation Methods and Implications for Drug Screening. ACS Biomater. Sci. Eng. 4, 410–420 (2018).

55. Benson, T. O., Islam, M. A., Liu, K. & Versypt, A. N. F. Agent-based modeling reveals impacts of cell adhesion and matrix remodeling on cancer collective cell migration phenotypes. bioRxiv 2024.12.23.630172 (2024) doi:10.1101/2024.12.23.630172.

56. Mungai, R. W., Hartman, R. J., Jolin, G. E., Piskorowski, K. W. & Billiar, K. L. Towards a more objective and high-throughput spheroid invasion assay quantification method. Scientific Reports 2024 14:1 14, 1–13 (2024).

57. Ho, C. Z. et al. Protocol for AI-assisted quantitative analysis and setup of tumor spheroid invasion into tissue. STAR Protoc. 6, 104140 (2025).

58. Knode, B. K. et al. Adipose-mimetic granular hydrogels uncover biophysical cues driving breast cancer invasion. bioRxiv 2025.10.23.684224 (2025) doi:10.1101/2025.10.23.684224.

59. Shimpi, A. A. et al. Convergent Approaches to Delineate the Metabolic Regulation of Tumor Invasion by Hyaluronic Acid Biosynthesis. Adv. Healthc. Mater. 12, (2023).

60. Mosier, J. A., Fabiano, E. D., Ludolph, C. M., White, A. E. & Reinhart-King, C. A. Confinement primes cells for faster migration by polarizing active mitochondria. Nanoscale Adv. 6, 209–220 (2023).

61. Keys, J., Cheung, B. C. H., Elpers, M. A., Wu, M. & Lammerding, J. Rear cortex contraction aids in nuclear transit during confined migration by increasing pressure in the cell posterior. J. Cell Sci. 137, (2024).

62. Diaz, G., Zuccarelli, A., Pelligra, I. & Ghiani, A. Elliptic Fourier analysis of cell and nuclear shapes. Computers and Biomedical Research 22, 405–414 (1989).

63. Kim, S. H. et al. Computer-aided image analysis of focal hepatic lesions in ultrasonography: Preliminary results. Abdom. Imaging 34, 183–191 (2009).

64. Ronneberger, O., Fischer, P. & Brox, T. U-Net: Convolutional Networks for Biomedical Image Segmentation. Lecture Notes in Computer Science (including subseries Lecture Notes in Artificial Intelligence and Lecture Notes in Bioinformatics) 9351, 234–241 (2015).

65. Berg, S. et al. ilastik: interactive machine learning for (bio)image analysis. Nature Methods 2019 16:12 16, 1226–1232 (2019).

66. Moriconi, C. et al. INSIDIA: A FIJI Macro Delivering High-Throughput and High-Content Spheroid Invasion Analysis. Biotechnol. J. 12, 1700140 (2017).

67. Akshay, A. et al. SpheroScan: a user-friendly deep learning tool for spheroid image analysis. Gigascience 12, 1–9 (2022).

68. Lacalle, D. et al. SpheroidJ: An Open-Source Set of Tools for Spheroid Segmentation. Comput. Methods Programs Biomed. 200, 105837 (2021).

69. Barrow, R. et al. SpheroidAnalyseR—an online platform for analyzing data from 3D spheroids or organoids grown in 96-well plates. J. Biol. Methods 9, e163 (2022).

70. Ilina, O., Bakker, G. J., Vasaturo, A., Hofmann, R. M. & Friedl, P. Two-photon laser-generated microtracks in 3D collagen lattices: principles of MMP-dependent and -independent collective cancer cell invasion. Phys. Biol. 8, 015010 (2011).

71. Ewald, A. J., Brenot, A., Duong, M., Chan, B. S. & Werb, Z. Collective Epithelial Migration and Cell Rearrangements Drive Mammary Branching Morphogenesis. Dev. Cell 14, 570–581 (2008).

72. Zanotelli, M. R. et al. Energetic costs regulated by cell mechanics and confinement are predictive of migration path during decision-making. Nature Communications 2019 10:1 10, 4185- (2019).

73. Ling, L. et al. Assessing cellular metabolic dynamics with NAD(P)H fluorescence polarization imaging. bioRxiv 2025.07.28.667273 (2025) doi:10.1101/2025.07.28.667273.

74. Mahdy, D. et al. AI-Powered Framework for Evaluating Drug Efficacy for Three-Dimensional In Vitro Cancer Models in Robot-Assisted Production. Advanced Robotics Research e202500049 (2026) doi:10.1002/ADRR.202500049.

75. Flesken-Nikitin, A. et al. Cell state dynamics during early stages of serous endometrial carcinoma. J. Pathol. 267, 289–303 (2025).

76. Flesken-Nikitin, A. et al. Pre-ciliated tubal epithelial cells are prone to initiation of high-grade serous ovarian carcinoma. Nature Communications 2024 15:1 15, 1–16 (2024).

