## Supplemental Figures 1-6 for "Shape Factor Analysis as a Quantitative Framework for Assessing Spheroid and Organoid Morphology and Invasiveness"

**A**

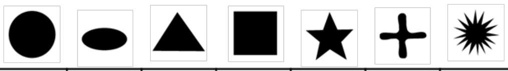

| | Image Resolution (pixels) | Circle | Oval | Triangle | Square | Star | X | Burst |
|---|---|---|---|---|---|---|---|---|
| Number of perimeter points in an interpolated ROI | 256 | 668 | 501 | 667 | 801 | 821 | 776 | 1951 |
| | 512 | 1336 | 1002 | 1334 | 1601 | 1645 | 1552 | 3932 |
| | 1024 | 2671 | 2004 | 2669 | 3201 | 3295 | 3103 | 7888 |
| Roundness | 256 | 1 | 0.48 | 0.86 | 1 | 0.95 | 0.98 | 0.99 |
| | 512 | 1 | 0.48 | 0.86 | 1 | 0.94 | 0.98 | 0.98 |
| | 1024 | 1 | 0.48 | 0.86 | 1 | 0.94 | 0.98 | 0.98 |
| Circularity | 256 | 1 | 0.83 | 0.6 | 0.79 | 0.33 | 0.25 | 0.05 |
| | 512 | 1 | 0.83 | 0.6 | 0.79 | 0.33 | 0.25 | 0.05 |
| | 1024 | 1 | 0.83 | 0.6 | 0.79 | 0.33 | 0.25 | 0.05 |
| Solidity | 256 | 1 | 1 | 1 | 1 | 0.54 | 0.48 | 0.44 |
| | 512 | 1 | 1 | 1 | 1 | 0.53 | 0.48 | 0.44 |
| | 1024 | 1 | 1 | 1 | 1 | 0.53 | 0.48 | 0.44 |
| Convexity | 256 | 1 | 1 | 1.002 | 0.999 | 0.848 | 0.782 | 0.344 |
| | 512 | 1 | 1 | 1.001 | 0.999 | 0.844 | 0.783 | 0.339 |
| | 1024 | 1 | 1 | 1 | 1 | 0.842 | 0.782 | 0.337 |
| Aspect Ratio | 256 | 1 | 2.07 | 1.17 | 1 | 1.06 | 1.03 | 1.02 |
| | 512 | 1 | 2.06 | 1.16 | 1 | 1.06 | 1.02 | 1.02 |
| | 1024 | 1 | 2.06 | 1.16 | 1 | 1.06 | 1.02 | 1.02 |

**B**

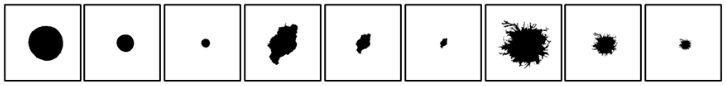


| Perimeter points | 1121 | 561 | 283 | 1573 | 771 | 373 | 7159 | 3579 | 1171 |
|---|---|---|---|---|---|---|---|---|---|
| Roundness | 0.97 | 0.97 | 0.97 | 0.57 | 0.57 | 0.57 | 0.9 | 0.9 | 0.9 |
| Circularity | 0.78 | 0.85 | 0.89 | 0.33 | 0.39 | 0.46 | 0.03 | 0.03 | 0.03 |
| Solidity | 0.98 | 0.97 | 0.96 | 0.84 | 0.84 | 0.83 | 0.64 | 0.64 | 0.64 |
| Convexity | 0.9 | 0.94 | 0.96 | 0.68 | 0.74 | 0.81 | 0.22 | 0.24 | 0.23 |
| Aspect Ratio | 1.03 | 1.03 | 1.03 | 1.75 | 1.76 | 1.76 | 1.12 | 1.11 | 1.12 |

**Supplemental Figure 1: Shape descriptor image resolution sensitivity analysis of the basic shape digital phantoms**. **A)** Shapes were generated in a 512x512 resolution image and scaled by 0.5 and 2 to proportionally fill 265x256 and 1024x1024 images respectively. ROIs were interpolated at a 1-pixel interval at each resolution with the number of perimeter pixels recorded at each size. Shape descriptors calculated at each resolution demonstrate independence from the image resolution. **B)** The sample spheroids in figure 1 were acquired and segmented from a 512 x 512 resolution image (1x). Spheroid ROIs were scaled at 0.5 and 0.25X to decrease the size and reduce the number of perimeter points in the ROI. Results show minimal variances in the shape factors as a function of the size of the ROI.

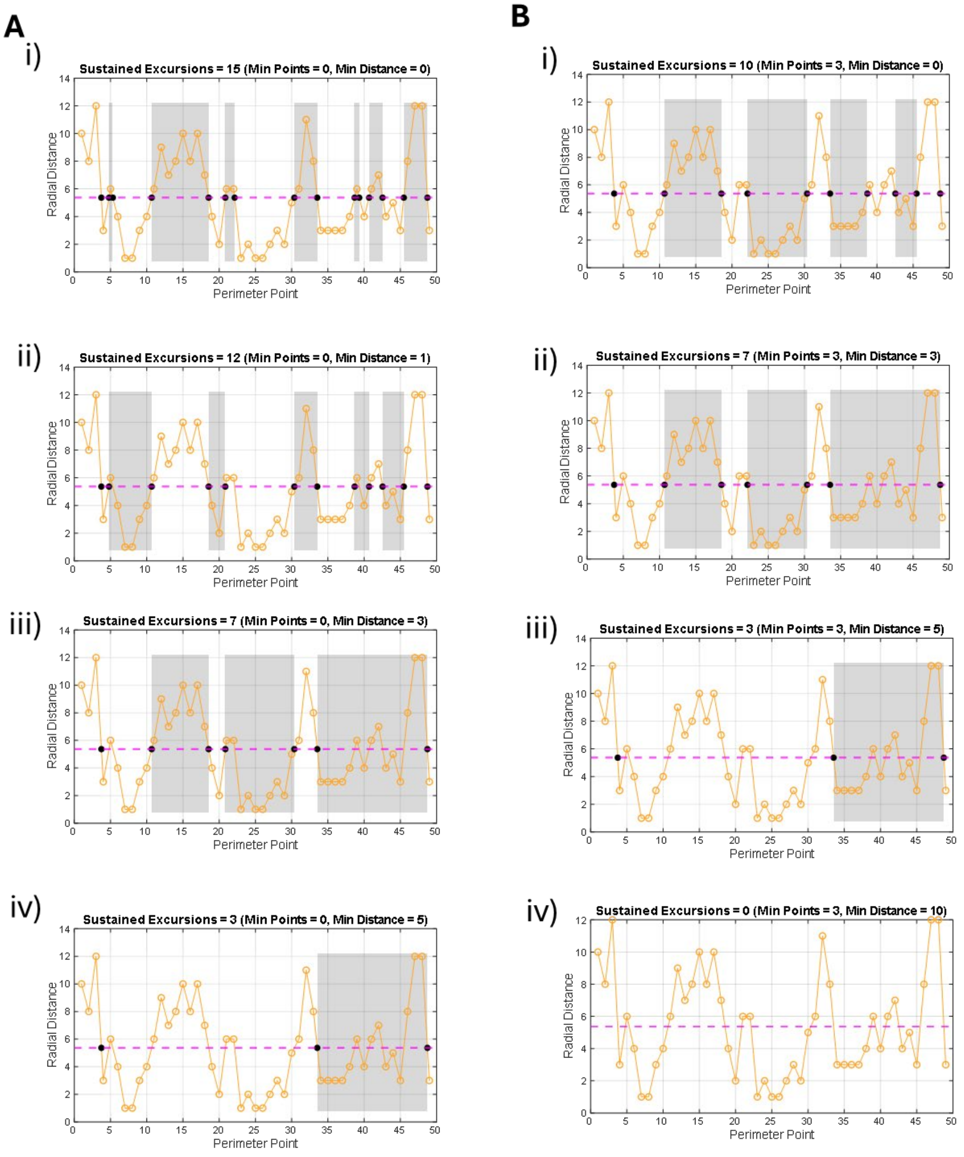


**Supplemental Figure 2:** Sample data to illustrate how altering the parameters of the average radial length crossing quantification (excursion length (minimum number of consecutive points to fall to one side of the average) and minimum distance) changes the detection of crossings and must be optimized to the relative amount of noise in a given data set. **A and B)** Sample vector plot with the radial lengths denoted by orange open circles and the average radial length plotted in magenta. Average Radial length crossings are denoted by black points. X-axis is the perimeter point index and the y-axis is the radial length value. **A**) With no minimum excursion length, the detected crossings are determined exclusively by the minimum crossing distance **i)** with no minimum distance counts there are 15 detected crossings. **ii)** 1-pixel minimum crossing distance yields 12 crossing **iii)** the minimum crossing distance to 3 points on either side of the axis reduces the number of detected crossings to 7 **iv)** 5-pixel minimum crossing distance gives 3 ARLC **Bi-iv)** An excursion length of 3 for all conditions yields 10, 7, 3, and 0 crossings for minimum crossing distances of 0, 3, 5, and 10 respectfully.

**A**

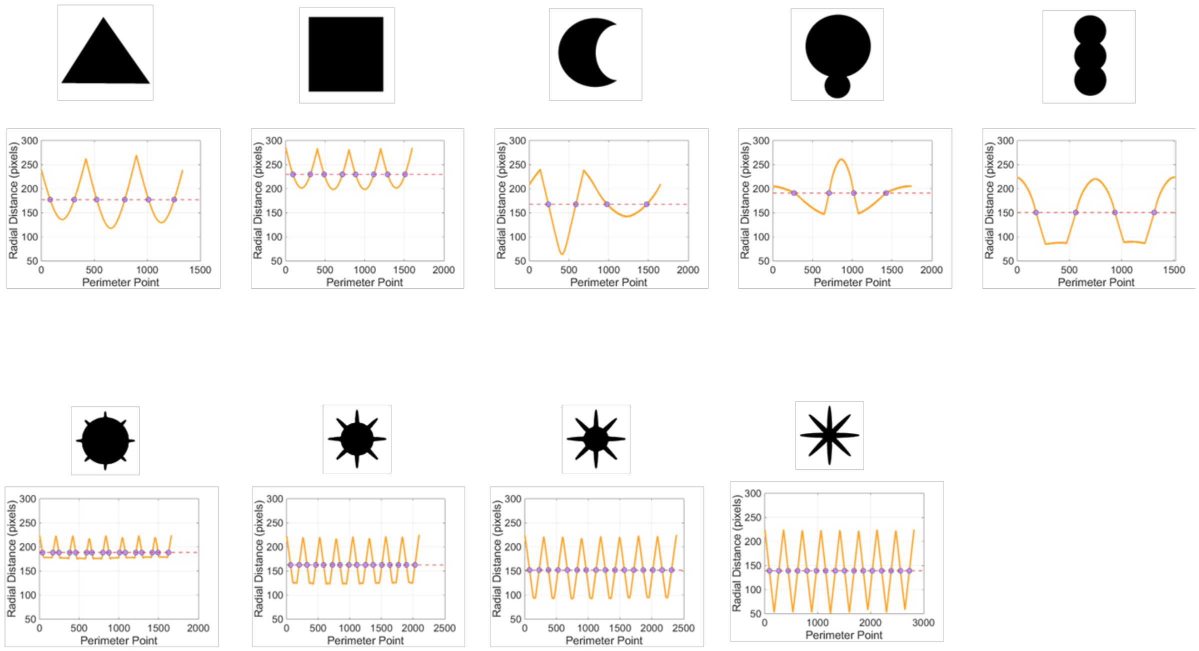


**B**

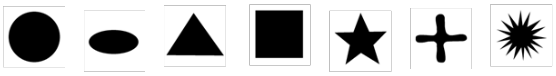

| | Pixel Resolution | Circle | Oval | Triangle | Square | Star | X | Burst |
|---|---|---|---|---|---|---|---|---|
| Average Radial Length Crossings | **256** | 2 | 4 | 6 | 8 | 10 | 8 | 32 |
| | **512** | 2 | 4 | 6 | 8 | 10 | 8 | 32 |
| | **1024** | 2 | 4 | 6 | 8 | 10 | 8 | 32 |
| Standard Deviation of Radial Length | **256** | 1 | 18 | 21 | 13 | 20 | 26 | 17 |
| | **512** | 1 | 40 | 41 | 25 | 40 | 50 | 34 |
| | **1024** | 1 | 74 | 82 | 51 | 79 | 102 | 70 |

**Supplemental Figure 3: Digital phantoms evaluated by shape descriptors and radial length analysis**. **A**) average radial length crossing plots for the digital phantoms with radial lengths in orange, the average in magenta, and crossings denoted by purple points. **B)** With ARLC detection parameters of a minimum excursion length of 3 and minimum crossing distance of 1, the resolution and corresponding change in the number of perimeter points did not impact the number of ARLC, however, the $SD_{RL}$ increased proportionally with the pixel area of the shape.

**A**

| Excursion length | 3 | | | 10 | | |
|---|---|---|---|---|---|---|
| Minimum crossing distance | **1** | **5** | **10** | **1** | **5** | **10** |
| ARLC 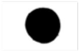 | 21 | 2 | 0 | 14 | 2 | 0 |
| ARLC 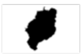 | 9 | 6 | 5 | 7 | 6 | 5 |
| ARLC 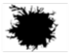 | 67 | 53 | 38 | 61 | 53 | 38 |

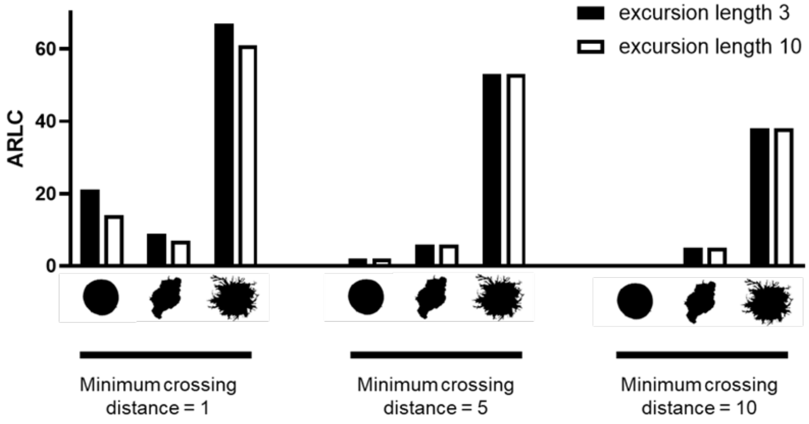


**B**

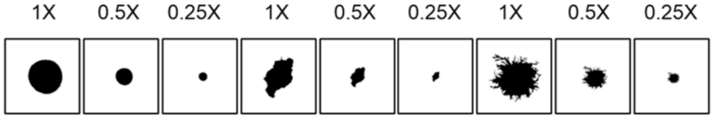


| | | | | | | | | | |
|---|---|---|---|---|---|---|---|---|---|
| **Perimeter points** | 1121 | 561 | 283 | 1573 | 771 | 373 | 7159 | 3579 | 1171 |
| **ARLC** | 2 | 1 | 0 | 6 | 5 | 4 | 53 | 38 | 24 |
| **$SD_{RL}$ (pixels)** | 2.7 | 1.9 | 1.6 | 23.3 | 11.7 | 5.9 | 23.4 | 11.0 | 6.1 |
| **Normalized $SD_{RL}$ (µm)** | 4.48 | 6.3 | 10.6 | 38.7 | 38.8 | 39.6 | 64.8 | 60.9 | 67.5 |

**Supplemental Figure 4: The impact of detection parameters and resolution of the sample spheroids on ARLC and $SD_{RL}$.** The average radial length crossing analysis was run on each spheroid sample with the following parameters: Excursion length of 3 with a minimum distance of 1, 5, or 10 pixels, and excursion length of 10 with a minimum distance of 1, 5, or 10 pixels. The average radial length crossings for each shape are displayed in the table. The orange highlighted values indicate the parameters used in the analysis in figure 3. **B)** Scaled spheroids using the excursion length of 3 and minimum crossing distance of 5 show that ARLC decreases with decreasing scale of the ROI in an image at a constant pixel resolution (512x 512 here).

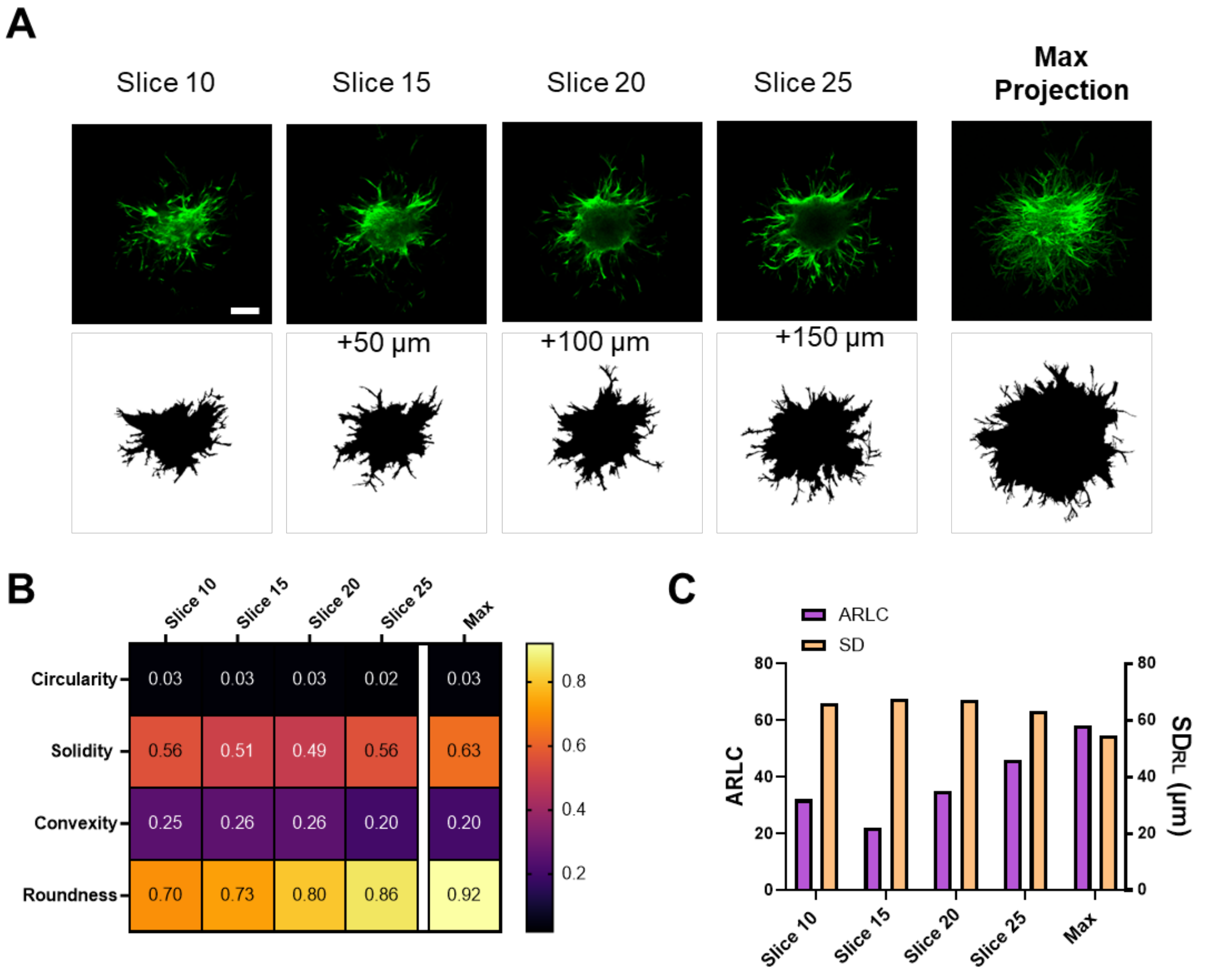


**Supplemental figure 5: Depth dependent shape factor analysis of optical slices of a spheroid acquired via confocal imaging**. **A)** Selected confocal slices of an F-actin-stained spheroid (green, 10 μm slices, 200 μm scale bar) and the corresponding thresholded and auto segmented (triangle threshold filter) ROIs displayed below. The maximum projection of the confocal stack corresponds to a projection of the top half of the spheroids from the top to the equatorial plane. **B)** Shape descriptor analysis of each slice and the corresponding max projection. **C)** Radial distance analysis reveals variations between slices with the highest ARLC count occurring in the max projection.

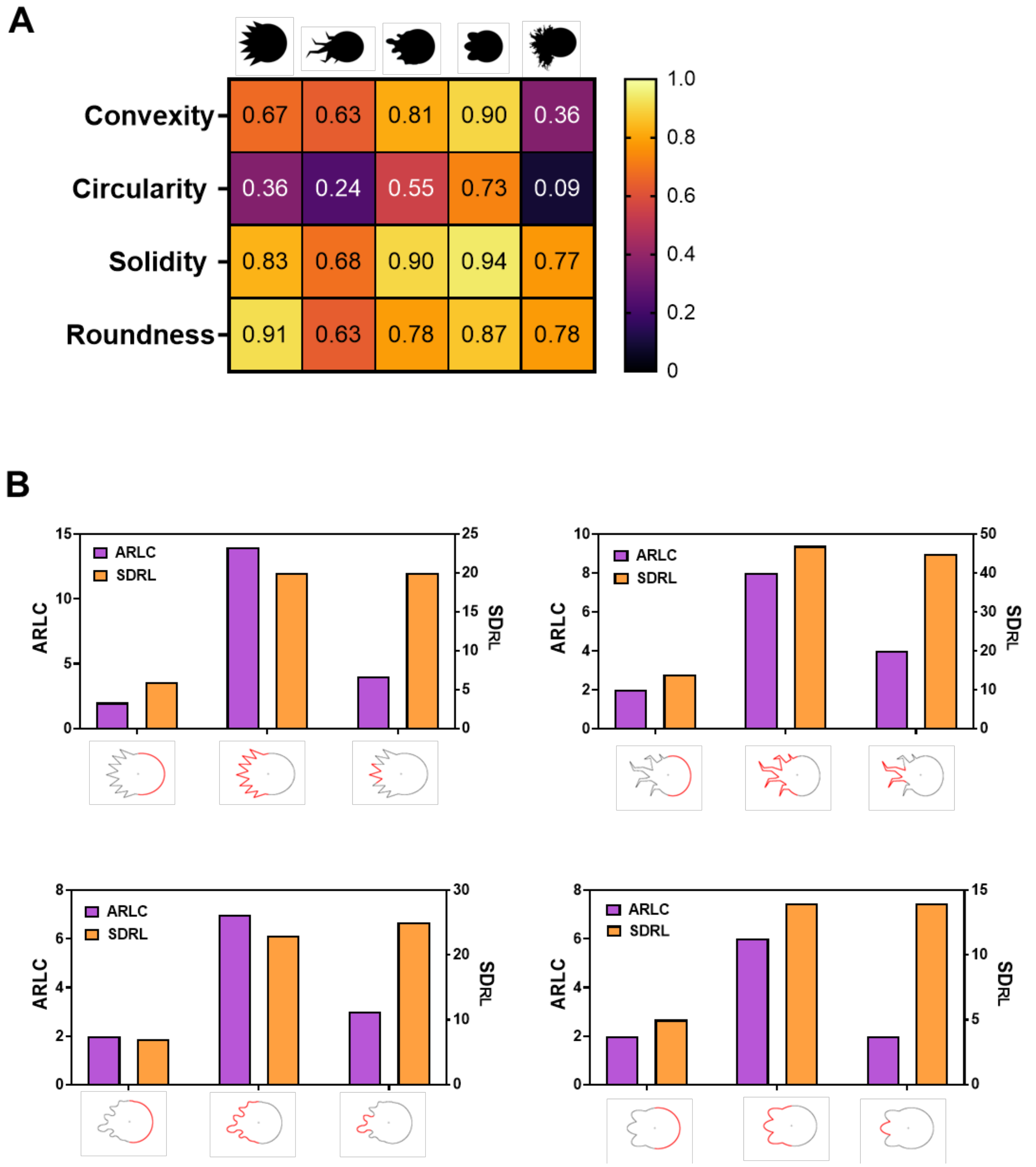


**Supplemental Figure 6: Analysis of asymmetric digital phantoms with simulated directed invasion to the left. A)** FIJI shape descriptor analysis of representative digital phantoms provides overall bulk analysis of the shapes with the limitation of not quantifying definable regions. **B)** Bar plots of the ARLC (purple), and $SD_{RL}$ (orange) of phantom segments of the regions 270-90°, 90-270°, and 157.5-202.5° respectively from left to right in each plot. The analyzed segments are highlighted in red on the phantom perimeters displayed below each plot.
